# LARP4 is a B cell-specific metabolic checkpoint for plasma cell differentiation and a therapeutic target in systemic lupus erythematosus

**DOI:** 10.64898/2026.07.10.737704

**Authors:** Huanzi Dai, Mingyang Zhang, Chun Lan, Fei Xiao, Junhui Deng, Hui Dong, Chao Han, Jian Zhou, Shufeng Wang, Jingxue Wang, Yaxing Hao, Yiwei Zhang, Zhaohui Zhang, Yi Sun, Jie Luo, Jun Zhu, Jingbo Zhang, Tingting Zhao, Xiangmei Chen, Yuzhang Wu, Di Yang, Yi Tian

## Abstract

RNA-binding protein LARP4 plays an important role in T cell activation and differentiation, but its role in B cell biology and the pathogenesis of systemic lupus erythematosus (SLE) remains unclear. This study found that LARP4 was specifically highly expressed in B cells of SLE patients and was positively correlated with disease activity. By constructing T cell-specific and B cell-specific conditional knockout mice, we found that deletion of LARP4 in B cells, but not in T cells, significantly alleviated pristane-induced and Bm12-induced lupus nephritis. Further analysis showed that LARP4 deletion selectively inhibited B cell differentiation into plasma cells, but did not affect germinal center B cell formation. Integrated transcriptomic and metabolomics analyses revealed that this effect is due to reduced phosphatidic acid synthesis and decreased mTORC1 activity caused by mitochondrial oxidative phosphorylation dysfunction. Furthermore, we used LIPEP, a LARP4 inhibitory peptide that effectively mimicked the therapeutic effects of LARP4 gene knockout in the MRL/lpr spontaneous lupus model and outperformed cyclophosphamide in reducing glomerular immune complex deposition and improving extrarenal dermatitis. These results indicates that LARP4 is a key metabolic checkpoint regulating B cell differentiation into Plasma cells and suggest that it may be a potential therapeutic target for SLE.

## Introduction

Systemic lupus erythematosus (SLE) is a chronic autoimmune disease characterized by B-cell hyperactivation and the production of pathogenic autoantibodies, resulting in multi-organ damage, particularly lupus nephritis ^1–3^. B cells play a central role in the pathogenesis of SLE, exerting functions that extend beyond autoantibody secretion to encompass antigen presentation and cytokine production ^4–6^. The terminal differentiation of B cells into antibody-secreting plasma cells is a critical step in the production of pathogenic autoantibodies, a process accompanied by profound metabolic remodeling ^7,8^.

The differentiation of B cells into plasma cells imposes extremely high demands on their energy and biosynthetic capacity. Resting and germinal center (GC) B cells primarily rely on glycolysis to meet their metabolic needs, whereas plasma cells markedly upregulate oxidative phosphorylation (OXPHOS) and mitochondrial respiration to support the robust synthesis and secretion of immunoglobulins ^9–11^. This metabolic switch is crucial for the proper execution of the plasma cell differentiation program. Studies have shown that mitochondrial DNA in B lymphocytes regulates the flux of the tricarboxylic acid (TCA) cycle by maintaining oxidative phosphorylation, thereby supporting humoral immunity and plasma cell maturation^12,13^. The integrity of the mitochondrial respiratory chain is intrinsically coupled to the efficient functioning of the TCA cycle: mitochondrial DNA encodes key subunits of respiratory chain complexes I, III, IV, and V, and the fidelity of its replication and transcription is indispensable for maintaining OXPHOS function^14,15^. When this metabolic switch is disrupted, both plasma cell formation and antibody production are severely impaired ^16,17^. However, the molecular mechanisms regulating this critical metabolic checkpoint remain incompletely understood.

In recent years, multiple studies have revealed key factors regulating B cell metabolism and differentiation. Monocarboxylic acid transporter 1 (MCT1) ensures appropriate intracellular pyruvate concentration, maintaining H3K27 acetylation levels to promote AID transcription, thereby influencing IgG antibody production. MCT1 deficiency can significantly reprogram glucose metabolism in B cells, shifting from glycolysis to oxidative phosphorylation, ultimately impairing antibody class switch recombination (CSR)^18^. Thioredoxin (Trx) acts as a metabolic rheostat for regulatory B cells (Bregs), controlling OXPHOS-dependent Breg differentiation. Breg cells highly express TXN (encoding Trx) with downregulated TXNIP. inhibiting Trx disrupts mitochondrial membrane potential and impairs their immunosuppressive function. In systemic lupus erythematosus (SLE) patients, Breg numbers and function are reduced, accompanied by mitochondrial abnormalities and Trx deficiency, and exogenous Trx can restore Breg numbers and function^19^. Furthermore, the mitochondrial iron transporter ABCB7 has been identified as a critical factor for mouse B cell development, proliferation, and CSR; its deletion leads to severe pro-B stage blockade, intracellular iron accumulation, and replication-associated DNA damage, but does not induce ROS or ferroptosis^20^. IL-2 signaling directly activates the mTOR-AKT-Blimp-1 axis in B cells, driving early metabolic reprogramming and promoting extrafollicular plasma cell differentiation, which determines the trajectory of early humoral immune responses^21^. Activation of pyruvate kinase M2 (PKM2) has been found to reprogram mitochondrial metabolism in CD8^+^T cells, enhancing effector function by influencing one-carbon metabolism and DNA methylation ^22^. Additionally, the transcription factor ZEB2 has been identified as a key driver of age-associated B cell (ABC) generation, directly regulating ABC differentiation and pathogenic function through the JAK-STAT signaling pathway^23,24^. ABCs in autoimmune diseases such as lupus are highly dependent on glycolysis to meet their functional demands, and ZEB2 deletion can alleviate autoimmune pathology^8^. These studies collectively indicate that different immune cells possess precise metabolic checkpoints during differentiation, and abnormalities in these checkpoints are closely associated with the onset and progression of autoimmune diseases.

However, while most known metabolic regulators described above are metabolic enzymes, transporters, or signaling pathway molecules, the role of RNA-binding proteins (RBPs) in the metabolic reprogramming of immune cells, particularly in B cells, remains poorly understood. Recent studies using CRISPR/Cas9 functional screening have identified that the Csde1-Strap complex regulates plasma cell differentiation by coupling the translation and degradation of Bach2 mRNA^25^, suggesting that RBPs may be overlooked key regulators in B cell fate determination. As a canonical RNA-binding protein, La-related protein 4 (LARP4) has been extensively characterized in T cells. In CD4⁺ T cells, LARP4 promotes cell activation and regulates their differentiation into Th1, Th2, and Th17 subsets ^26,27^. Recent studies indicate that LARP4 supports mitochondrial respiratory function by enhancing the translation efficiency of nuclear-encoded OXPHOS subunit mRNA^28,29^. In a CD8^+^T cell exhaustion model, LARP4 drives mitochondrial dysfunction by specifically upregulating the translation of OXPHOS-related mRNAs, thereby contributing to the formation of the exhaustion phenotype ^28^. Given the research gap on RBPs in B cell metabolic regulation and the established core regulatory role of LARP4 in mitochondrial OXPHOS and cell differentiation in T cells, a key question arises: Does LARP4 exert similar or distinct metabolic regulatory functions in B cells? Is it involved in the pathogenesis of autoimmune diseases such as SLE? This remains unclear to date.

In this study, we found that LARP4 was specifically highly expressed in B cells of SLE patients and positively correlated with disease activity. Using T cell- and B cell-specific conditional knockout mice, we further demonstrated that, unlike in T cells, the deletion of LARP4 in B cells significantly alleviated disease progression in multiple lupus models. Mechanistically, LARP4 deficiency selectively blocked B-cell differentiation into plasma cells without affecting germinal center B-cell formation. This effect was attributed to reduced phosphatidic acid (PA) synthesis, which subsequently decreased mTORC1 activity as a result of mitochondrial OXPHOS dysfunction. Building on these findings, our previously designed LARP4 inhibitory peptide LIPEP effectively mimicked the therapeutic effect of gene knockout in MRL/lpr mice and outperformed the classic immunosuppressant cyclophosphamide in reducing glomerular deposits, inhibiting interstitial inflammation, and improving extrarenal dermatitis. These findings establish LARP4 as a B cell-specific metabolic checkpoint and highlight its potential as a target for SLE treatment.

## Results

### 1. LARP4 is highly expressed in SLE B cells and is associated with disease activity

To investigate whether LARP4 is involved in the pathogenesis of SLE, we first analyzed publicly available transcriptomic data of peripheral blood mononuclear cells (PBMCs) from SLE patients. Relative to healthy controls (HC), LARP4 mRNA levels in PBMCs from SLE patients were significantly elevated (Fig. 1a). To determine whether the increase in LARP4 was cell-type specific, we further analyzed its expression in different immune cell subsets. Notably, LARP4 expression levels in B cells from SLE patients were significantly higher than those in healthy controls, while no significant difference was observed in CD4^+^T cells (Fig. 1b). Furthermore, LARP4 expression levels in B cells of SLE patients correlated positively and significantly with the Systemic Lupus Erythematosus Disease Activity Index (SLEDAI) score (Fig. 1c). These data indicate that the aberrant expression of LARP4 in SLE is B-cell-specific and may contribute to disease progression. These findings prompted us to investigate the distinct functional roles of LARP4 in B cells versus T cells during SLE pathogenesis.

**Fig 1.**
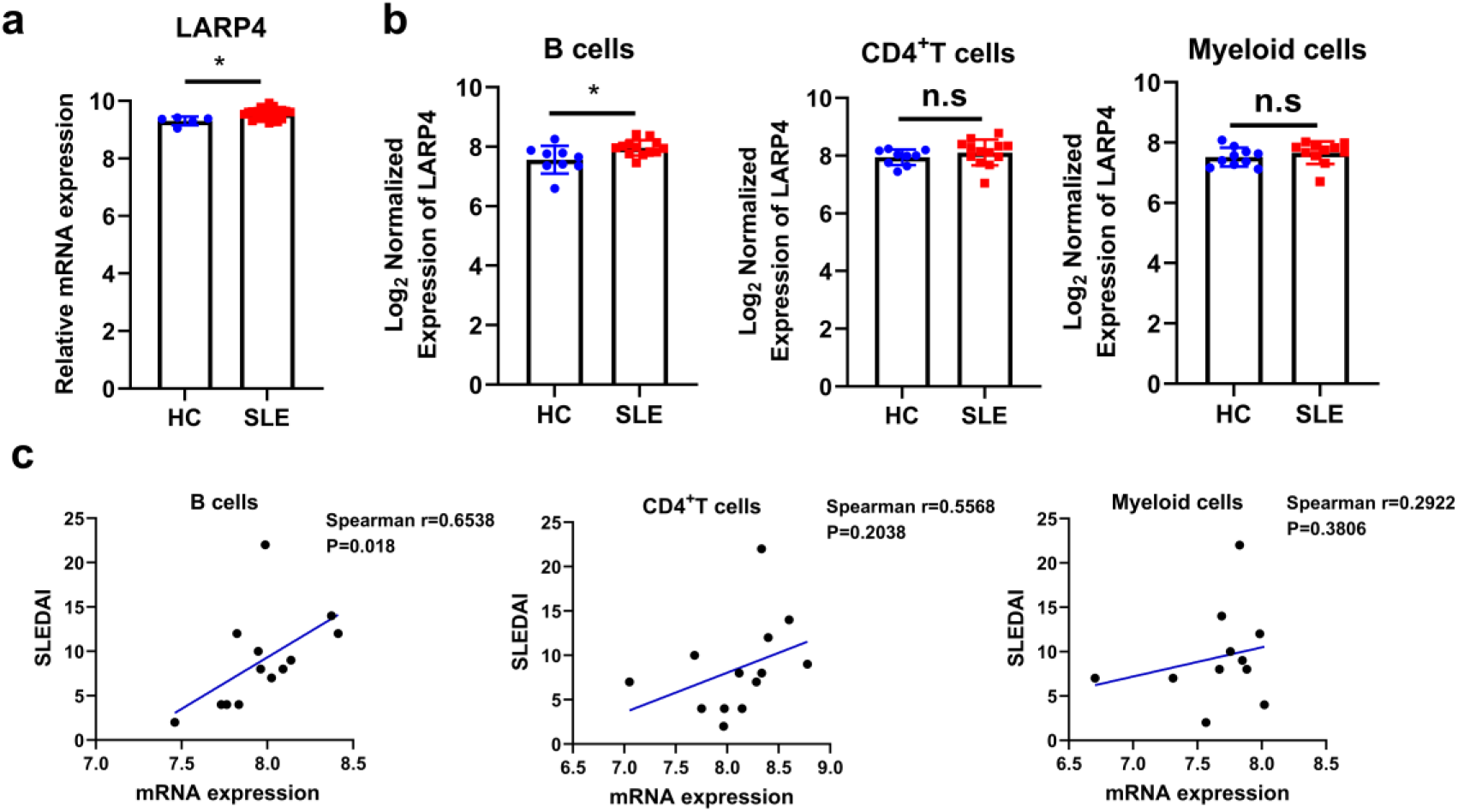
LARP4 expression is elevated in B cells from SLE patients and correlates with disease activity. **a.** LARP4 expression in PBMC of SLE patients and healthy controls, from data of GSE122459. **b.** MCT1 expression in B、CD4^+^T and myeloid cells of SLE patients and healthy controls, from data of GSE10325. **c.** Correlation between LARP4 mRNA expression in B CD4^+^T and myeloid cells from SLE patients and the SLEDAI score.

### 2. CD4^+^T cell LARP4 deficiency does not affect the severity of pristane-induced lupus

To clarify whether the expression of LARP4 in CD4⁺T cells is involved in the development and progression of SLE. we crossed Larp4-floxed mice with CD4-cre mice to generate CD4⁺T cell-specific Larp4 knockout mice (CD4-Larp4 CKO) and constructed a pristane-induced lupus model (Supplementary Fig. 1a). After 6 months, spleen size, spleen index, and splenocyte counts did not differ significantly between CD4-Larp4 CKO mice and Larp4^fl/fl^ mice (Supplementary Fig. 1b-d). Serological analysis showed no significant differences in anti-dsDNA IgG antibodies, total IgG levels, or 24-hour urine protein (Supplementary Fig. 1e–g). Renal PAS staining indicated that the two groups of mice exhibited comparable levels of glomerular endothelial cell proliferation, inflammatory cell infiltration, and renal tubulointerstitial injury (Supplementary Fig. 1h), and there was no significant reduction in the deposition of IgG and C3 in the glomeruli of CD4-Larp4 CKO mice (Supplementary Fig. 1i, j). Furthermore, H&E staining of lung tissue showed that pristane-induced diffuse alveolar hemorrhage (DAH) was also unaffected by T cell-specific LARP4 deficiency (Supplementary Fig. 2a,b). Additionally, flow cytometry revealed no significant difference in the frequency of Tfh cells (CD4⁺CXCR5⁺PD-1⁺) in the spleens of the two groups of mice (Supplementary Fig. 2c,d). These results indicate that the deletion of LARP4 in CD4⁺T cells is insufficient to protect mice from pristane-induced lupus nephritis, suggesting that the pathogenic role of LARP4 in SLE may occur in B cells rather than T cells.

### 3. B cell-specific LARP4 deletion alleviates Bm12-induced lupus disease progression

Given the results from the previous section suggesting that the pathogenic role of LARP4 in lupus pathogenesis is not primarily mediated by T cells, we shifted our focus to investigating the function of LARP4 in B cells. Therefore, to determine the role of LARP4 in B cell function in vivo, we generated the Larp4^fl/fl^ mouse strain and generated B cell-specific Larp4 knockout mice by crossing Larp4^fl/fl^ mice with CD19-Cre mice. Results confirmed the absence of LARP4 protein in samples from LARP4 conditional knockout (CKO) mice (Supplementary Fig. 3a,b). No significant differences were observed in spleen morphology, weight, or B cell follicular patterns between Larp4fl/fl and CD19-Cre Larp4fl/fl mice (Supplementary Fig. 4a-e). We also observed that B cell lineage-specific knockout (KO) of LARP4 did not affect B cell development in the bone marrow, plasma cell formation, or homeostasis of peripheral lymphoid organs (Supplementary Fig. 5a-k). Furthermore, the levels of innate antibodies were similar in both types of mice (Supplementary Fig. 4f-h).

Therefore, we utilized CD19-Cre Larp4^fl/fl^ mice and to establish a Bm12-induced lupus model (Fig. 2a). In contrast to the lack of efficacy observed with T cell-specific knockout, B cell-specific deletion led to a significant and consistent alleviation of lupus-like disease. Compared to Larp4^fl/fl^ mice, B-Larp4 CKO mice showed reduced splenomegaly (Fig. 2b), with significantly decreased splenocyte counts, spleen weight, and spleen index (Supplementary Fig. 6a-c). Secondly, serological analysis revealed a significant reduction in pathogenic autoantibodies, with anti-dsDNA IgG and ANA IgG levels in the serum of B-Larp4 CKO mice being significantly lower than those in Larp4fl/fl mice (Fig. 2c-f). Notably, total IgM levels were not significantly reduced and exhibited a non-significant upward trend (Supplementary Fig. 6d-f). Finally, renal pathology was markedly ameliorated. Histological examination with HE staining showed that endocapillary proliferation, neutrophil infiltration, ce-lular crescent, and tubular interstitial inflammation. were significantly reduced in B-Larp4 CKO mice (Fig. 2g). Correspondingly, IgG and C3 deposition in the glomeruli were also significantly reduced (Fig. 2h, i), while IgM deposition remained largely unaffected (Supplementary Fig. 6g, h). These data collectively indicate effective alleviation of lupus nephritis.

**Fig 2.**
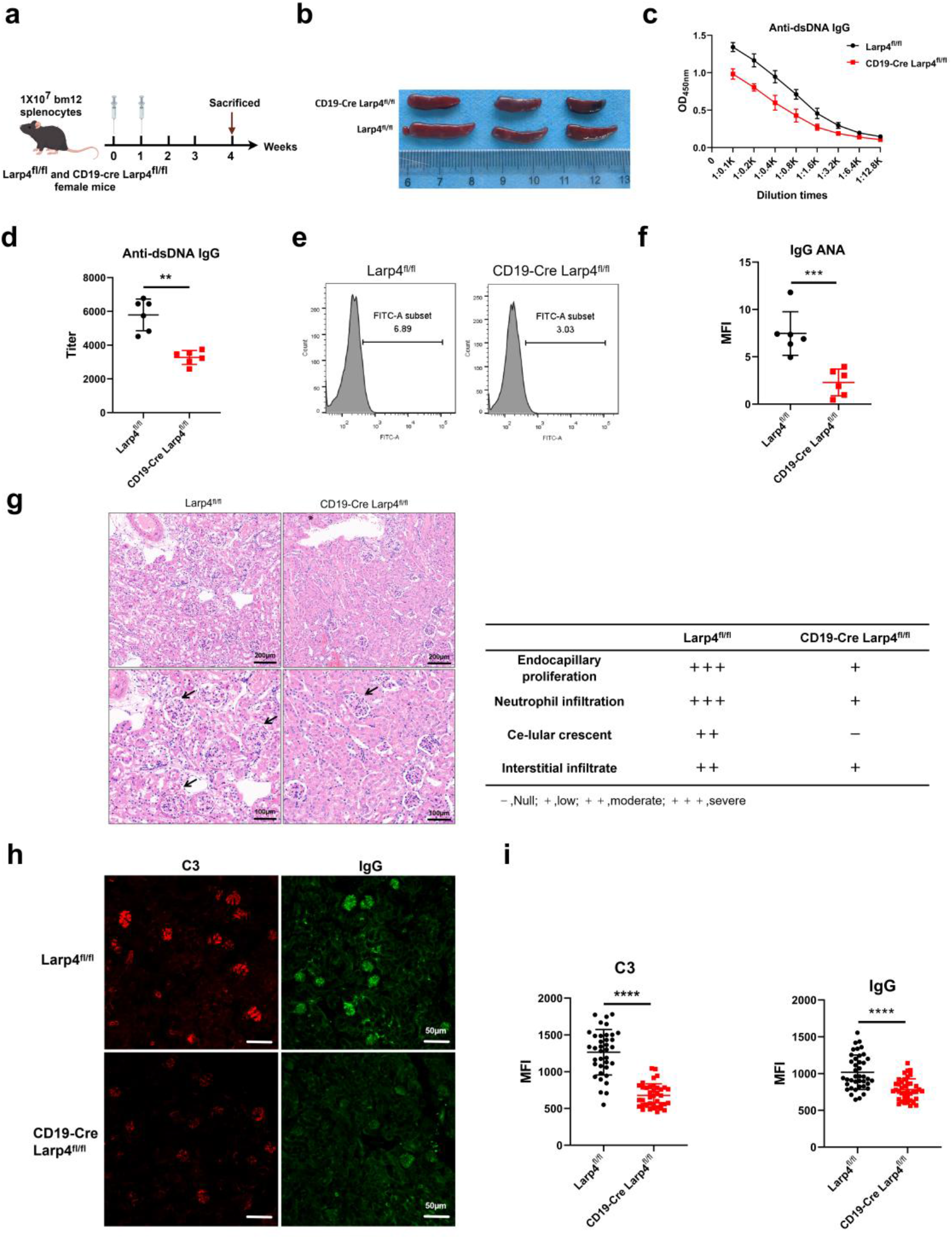
B cell-specific LARP4 deficiency attenuates Bm12-induced lupus nephritis. **a.** Schematic representation of the experimental design. Bm12 CD4⁺T cells were adoptively transferred into B-Larp4 CKO (CD19-Cre Larp4^fl/fl^) and Larp4^fl/fl^ mice. **b.** Representative photographs of spleens from Larp4^fl/fl^ and B-Larp4 CKO mice in bm12-induced SLE model. **c,d**. OD450 nm of serum anti-dsDNA IgG and their titers measured by ELISA. **e,f.** Serum levels of anti-nuclear antibody (ANA) IgG measured by Flow cytometry. **g,** Representative H&E-stained kidney sections (left, scale bar=100μm) and glomerular histopathology scores (right), including endocapillary proliferation, neutrophil infiltration, ce-lular crescent, and tubular interstitial inflammation. **h,i.** Representative immunofluorescence staining of IgG and C3 deposition in kidney glomeruli (**h,** scale bar = 50 μm) and quantification of fluorescence intensity**(i)**. Data are presented as mean ± SEM. *p < 0.05, **p < 0.01, ***p < 0.001 by two-tailed Student’s t-test.

### 4. LARP4 deficiency inhibits B-cell differentiation into plasma cells, without significant effect on GCB cells

To investigate the cellular mechanisms underlying the attenuated lupus phenotype in B-Larp4 knockout mice, we analyzed germinal center B (GCB) cells and plasma cells in the spleens of B-Larp4 CKO and Larp4^fl/fl^ mice immunized to induce the Bm12 lupus model. Flow cytometry analysis revealed that the frequency of GCB cells (B220^+^GL-7^+^CD95^+^) did not differ significantly between B-Larp4 CKO and Larp4^fl/fl^ mice (Fig. 3a,b); however, the frequency of plasma cells (B220^+^CD138^+^) was significantly reduced in B-Larp4 CKO mice compared to Larp4^fl/fl^ mice (Fig. 3c, d). Notably, in addition to the reduction in plasma cells, the frequency of Tfh cells (CD4^+^CXCR5^+^PD-1^+^) in the spleens of B-Larp4 CKO mice was also significantly lower than that in Larp4fl/fl mice (Supplementary Fig. 7a, b).

**Fig 3.**
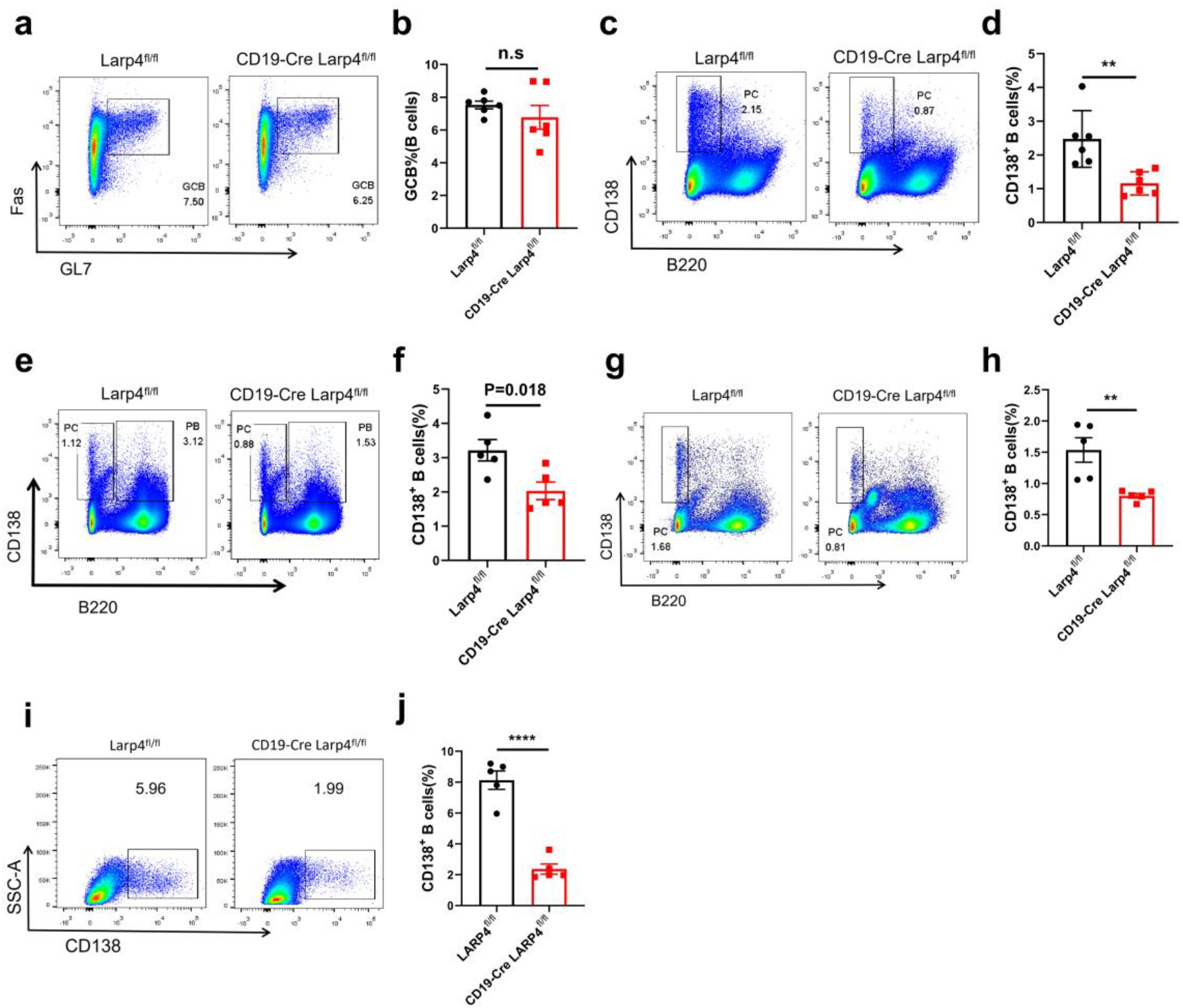
LARP4 deficiency impairs plasma cell differentiation without affecting germinal center B cell formation. **a,b.** Flow cytometry analysis of splenic GCBC from Larp4^fl/fl^ and B-Larp4 CKO mice in the bm12-induced SLE model**(a)** and the percentage of GCB cells is presented **(b)**. **c,d.** Flow cytometry analysis of plasma cells in the spleens of the same bm12-induced SLE model **(c)** and the percentage of plasma cells is presented **(d)**. **e,f.** Flow cytometry analysis of plasma cells in the spleens of Larp4^fl/fl^ and B-Larp4 CKO mice at day 28 with NP_33_-KLH immunization **(e)** and the percentage of plasma cells is presented **(f)**.**g,h.** Flow cytometry analysis of plasma cells in the spleens of Larp4^fl/fl^ and B-Larp4 CKO mice at day 14 with SRBC immunization **(g)** and the percentage of plasma cells is presented **(h)**.**i,j.** Flow cytometry analysis of CD138⁺B cells stimulated with LPS and IL-4 at day 3 from Larp4^fl/fl^ and B-Larp4 CKO mice **(i)** and the percentage of CD138⁺B cells is presented **(j)**. Data are presented as mean ± SEM. *p < 0.05, **p < 0.01, ***p < 0.001 by two-tailed Student’s t-test.

To further validate this observation, we immunized two mouse strains with T cell-dependent (TD) model antigens NP_33_-KLH and sheep red blood cells (SRBC), and quantified the differentiation of GCB cells and plasma cells via flow cytometry. The results showed that LARP4 deficiency consistently impaired plasma cell differentiation (Fig. 3e-h) but did not affect GCB cell formation (Supplementary Fig. 7c-f). Immunohistochemistry analysis using peanut agglutinin (PNA, a GCB cell marker) further revealed no significant differences in the number and size of GCB cells between B-Larp4 CKO mice and Larp4^fl/fl^ mice (Supplementary Fig. 7 g, right panel), nor in the frequency of Ki67-positive GCB cells (Supplementary Fig. 7 g, left panel). Finally, we validated these findings using an LPS+IL-4-induced in vitro plasma cell differentiation model. Flow cytometry analysis showed that LARP4 deficiency led to a significant reduction in the percentage of plasma cells (CD138^+^) (Fig. 3i, j). This result is consistent with findings from the Bm12 in vivo model, as well as the NP_33_-KLH and SRBC immunization models. These data indicate that LARP4 deficiency selectively inhibits plasma cell differentiation without affecting GCB cell formation.

### 5. LARP4 deficiency leads to mitochondrial dysfunction in plasma cells

To elucidate the molecular mechanism by which LARP4 deficiency selectively blocks plasma cell differentiation, we performed RNA sequencing and quantitative proteomic analyses on splenic plasma cells (B220⁻CD138⁺) from Bm12 lupus model mice. Transcriptomic analysis revealed that LARP4 deficiency led to 1,870 downregulated genes and 646 upregulated genes (Fig. 4a). KEGG enrichment analysis indicated that these downregulated genes were significantly enriched in pathways closely related to mitochondrial function, such as oxidative phosphorylation, reactive oxygen species and thermogenesis (Fig. 4b). Gene set enrichment analysis (GSEA) further confirmed a global downregulation of OXPHOS pathway genes in the B-Larp4 CKO group (Fig. 4c). Heatmap analysis revealed that in LARP4-deficient plasma cells, almost all nuclear-encoded OXPHOS subunit genes (Complex I–V) were consistently downregulated (Fig. 4d). Proteomic analysis results were highly consistent with the transcriptome, which revealed 494 downregulated proteins and 170 upregulated proteins (Fig. 4e). GSEA using MSigDB gene sets showed that the normalized enrichment score (NES) of oxidative phosphorylation-related pathways was significantly reduced in the CKO group (Fig. 4f). Consistent with this, heatmap visualization further revealed that the protein levels of nuclear-encoded OXPHOS subunits were globally downregulated in LARP4-deficient plasma cells (Fig. 4g).

**Fig 4.**
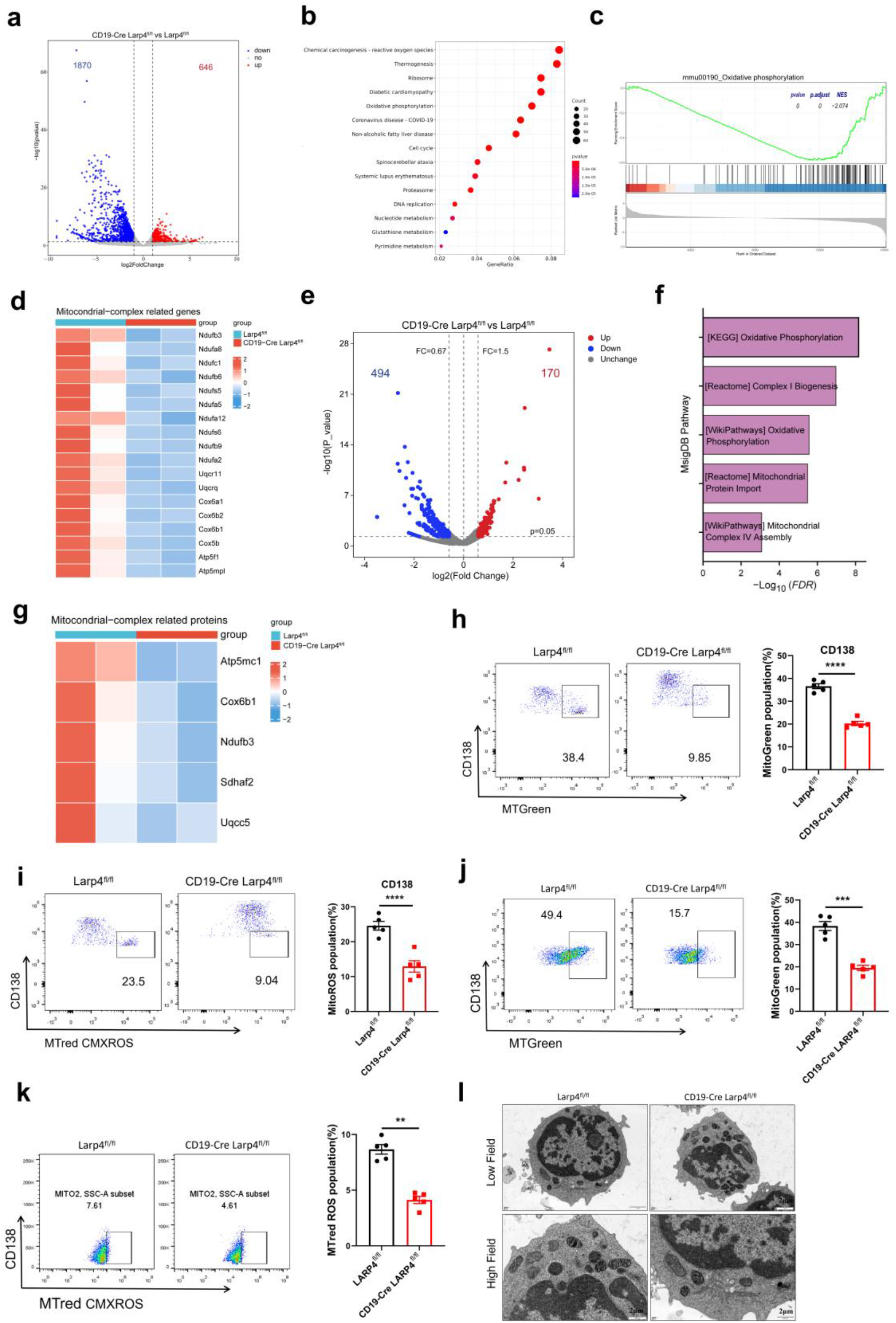
LARP4 deficiency causes mitochondrial dysfunction in plasma cells. **a.** Volcano plot showing differentially expressed genes (DEGs) identified by RNA-seq between B-Larp4 CKO and control plasma cells from Bm12 lupus mice. Red dots indicate significantly upregulated genes (n = 646), blue dots indicate significantly downregulated genes (n = 1,870). **b.**KEGG pathway enrichment analysis of the downregulated transcripts. **c.** Gene set enrichment analysis (GSEA) of the transcriptome data showing significant downregulation of the KEGG_OXIDATIVE_PHOSPHORYLATION gene set in B-Larp4 CKO cells. The normalized enrichment score (NES) and P value are indicated. **d.** Heatmap displaying the expression levels of nuclear-encoded OXPHOS subunit genes (Complex I–V) in B-Larp4 CKO and Larp4^fl/fl^ plasma cells. **e.** Volcano plot of differentially abundant proteins identified by quantitative proteomics between B-Larp4 CKO and Larp4^fl/fl^ plasma cells. Blue dots represent significantly downregulated proteins (n = 494), red dots represent significantly upregulated proteins (n = 170). **f.** GSEA performed on the proteome using MSigDB curated gene sets. The Oxidative phosphorylation-related gene sets show a significant decrease in normalized enrichment score (NES) in B-Larp4 CKO cells. **g.** Heatmap of the protein levels of nuclear-encoded OXPHOS subunits (Complex I–V). **h,i.** Flow cytometry analysis of mitochondrial membrane potential (MitoTracker Green staining, **h**) and mitochondrial mass (MitoTracker Red CMXRos staining, **i**) in splenic plasma cells from Larp4^fl/fl^ and B-Larp4 CKO mice in the Bm12-induced SLE model. **j,k.** Flow cytometry analysis of mitochondrial membrane potential (MitoTracker Green staining, **j**) and mitochondrial mass (MitoTracker Red CMXRos staining, **k**) in plasma cells stimulated with LPS and IL-4 for 3 days in vitro. **l.** Representative transmission electron microscopy (TEM) images of mitochondria in plasma cells from Larp4^fl/fl^ and B-Larp4 CKO mice. Data are presented as mean ± SEM.*p < 0.05, **p < 0.01, ***p < 0.001 **** p < 0.001 by two-tailed Student’s t-test.

The consistent results from the transcriptome and proteome prompted us to further validate mitochondrial functional phenotypes. Flow cytometry revealed that LARP4-deficient plasma cells exhibited a significant reduction in mitochondrial membrane potential, along with a marked decrease in mitochondrial mass (Fig. 4h, i), indicating severe impairment in both function and biomass. These functional defects were also validated in an independent in vitro LPS+IL-4 plasma cell differentiation system (Fig.4j,k). Transmission electron microscopy (TEM) provided direct ultrastructural evidence of mitochondrial injury: In contrast to the dense matrix and well-defined cristae observed in control plasma cells, LARP4-deficient plasma cells displayed reduced matrix density, sparse and fragmented cristae, vacuolization, and mitochondrial swelling (Fig. 4l).

In summary, these data collectively establish a robust mechanistic model: LARP4 deficiency leads to a global downregulation of nuclear-encoded OXPHOS subunits at both mRNA and protein levels, thereby inducing mitochondrial dysfunction. During the differentiation of B cells into plasma cells—a process characterized by surging energy demands—this dysfunction manifests as a lethal metabolic bottleneck, ultimately blocking terminal differentiation. This finding is highly consistent with LARP4’s role as a translational enhancer of nuclear-encoded OXPHOS mRNAs in T cells.

### 6. LARP4 deletion blocks plasma cell differentiation via the PA-mTORC1-Blimp-1 axis

Severe mitochondrial dysfunction was observed in LARP4-deficient plasma cells, suggesting potential compromise in downstream metabolic pathways crucial for cell signaling. To investigate this, we performed untargeted metabolomic analysis on flow cytometry-sorted splenic plasma cells from Bm12 lupus mice. Principal component analysis (PCA) revealed a clear metabolic separation between B-LARP4 CKO and control plasma cells (Fig. 5a). A total of 128 differentially abundant metabolites were identified, of which 53 were upregulated and 75 were downregulated (Fig. 5b). KEGG pathway enrichment analysis of the differential metabolites further revealed widespread metabolic perturbations. Among these, the valine, leucine, and isoleucine degradation pathway was significantly enriched, indicating impaired amino acid catabolism. The ubiquinone and other terpenoid-quinone biosynthesis pathway was also significantly affected, and the synthesis of ubiquinone (coenzyme Q), a key component of the electron transport chain, was hindered, directly limiting mitochondrial respiratory function (Fig. 5c). This indicates that LARP4 deficiency causes systemic disruption of the energy metabolism network in plasma cells. Notably, the metabonomics analysis revealed a series of metabolite changes highly consistent with mitochondrial dysfunction. Among these, the levels of L-arginine and thiamine (hydrochloride) were significantly reduced, while methylmalonic acid and ubiquinone-1 showed marked accumulation (Fig.5d). The accumulation of methylmalonic acid typically suggests impaired generation of succinyl-CoA in the TCA cycle, and the reduction of thiamine, a cofactor for key TCA cycle enzymes, further supports restricted flux through this cycle. The upregulation of ubiquinone-1 may reflect dysfunction or compensatory responses of the electron transport chain complexes. The coordinated changes in these metabolites collectively point to a core conclusion: impaired OXPHOS complex function leads to blockade and reprogramming of TCA cycle flux.

**Fig 5.**
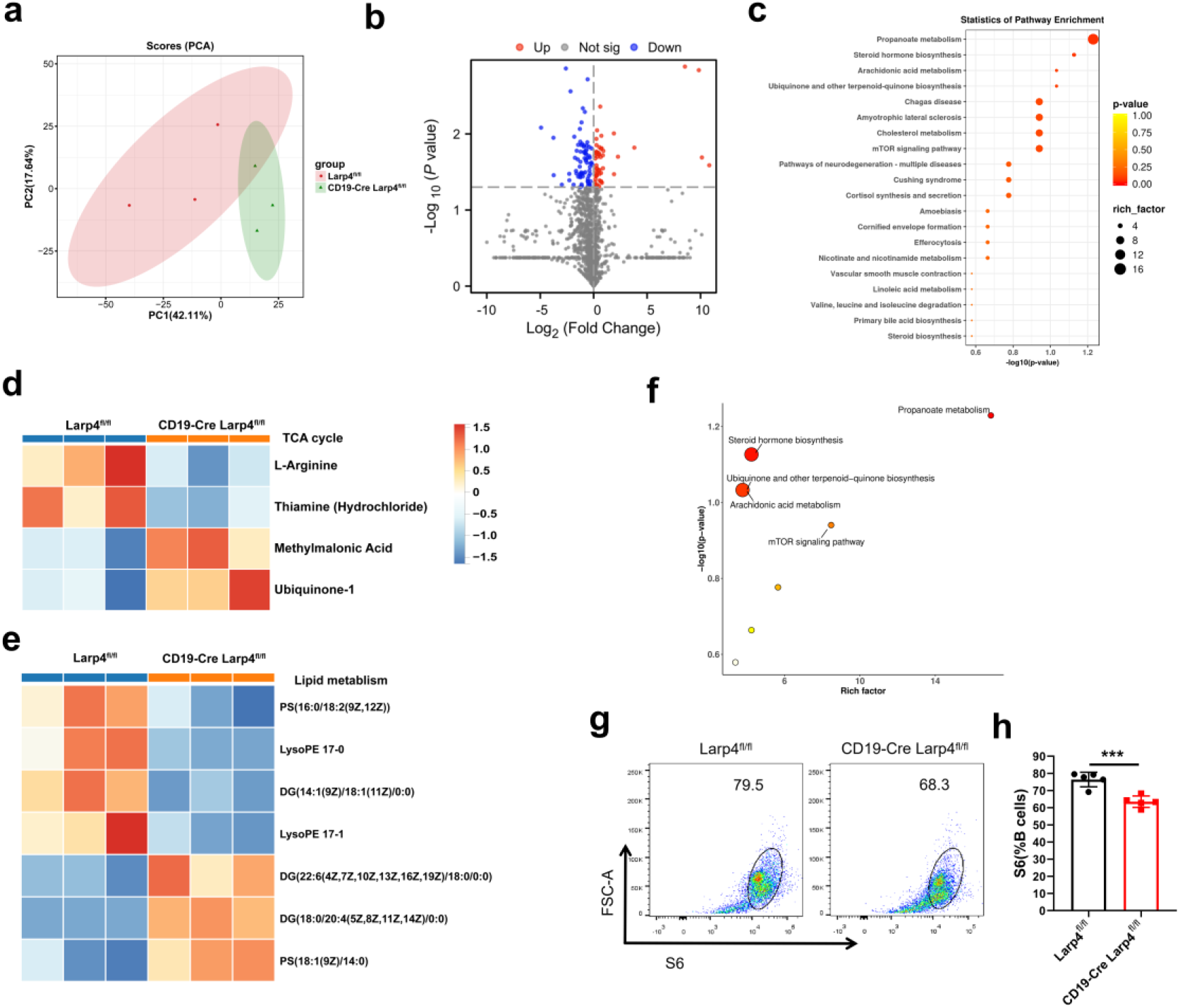
LARP4 deficiency disrupts TCA cycle flux and impairs PA-mTORC1 signaling in plasma cells. **a.** PCA of metabolic profiles in splenic plasma cells sorted from Larp4^fl/fl^ and B-Larp4 CKO mice in the Bm12-induced SLE model. **b.** Volcano plot showing differentially abundant metabolites. **c.** KEGG pathway enrichment analysis of differentially abundant metabolites. **d.** Heatmap showing metabolites in glycolytic pathway, TCA cycle in Larp4^fl/fl^ and B-Larp4 CKO plasma cells. **e.** Abundance of various phosphatidic acid (PA) species in Larp4^fl/fl^ and B-Larp4 CKO plasma cells. **f.** Pathway analysis of differential metabolites in Larp4^fl/fl^ and B-Larp4 CKO plasma cells. **g,h**. Flow cytometry analysis of pS6 levels in plasma cells from Larp4^fl/fl^ and B-Larp4 CKO mice in the Bm12-induced SLE model **(g)** and the percentage of pS6⁺ plasma cells is presented **(h)**, n = 4-5 biological replicates. Data are presented as mean ± SEM, ****p < 0.0001 by two-tailed Student’s t-test.

Importantly, this TCA cycle imbalance impaired de novo lipid biosynthesis, thereby reshaping the metabolic profile of glycerophospholipids (GPLs). We observed a divergent trend in different GPL precursors and derivatives: specifically, the levels of short-chain saturated/monounsaturated species, such as PS (16:0/18:2(9Z,12Z)), LysoPE 17-0, LysoPE 17-1, and DG (14:1(9Z)/18:1(11Z)/0:0), decreased, whereas the levels of long-chain polyunsaturated species, such as DG (22:6(4Z,7Z,10Z,13Z,16Z,19Z)/18:0/0:0) and DG (18:0/20:4(5Z,8Z,11Z,14Z)/0:0), increased (Fig. 5e). This pattern closely aligns with the lipid signature previously reported in a B-cell mitochondrial respiratory defect model: a reduction in short-chain saturated and monounsaturated acyl chains in GPLs, accompanied by an increase in long-chain polyunsaturated acyl chains^12^. As a core precursor for GPL biosynthesis (including PS and PE), phosphatidic acid (PA) availability may be diminished by impaired TCA cycle-driven de novo lipid synthesis. Therefore, the divergent regulation of the aforementioned GPL precursor metabolites implies that overall PA biosynthesis may be compromised. Furthermore, KEGG pathway analysis of differentially abundant metabolites also revealed significant enrichment of the mTOR signaling pathway, primarily driven by altered lipid species (Fig. 5f). Given that PA is a direct activator of mTORC1^31,32^, this result further supports the potential inhibition of the PA-mTORC1 axis.

To further assess mTORC1 activity via pS6 levels, flow cytometry analysis revealed significantly reduced pS6 levels in LARP4-deficient plasma cells in the Bm12 lupus mouse model (Fig. 5g, h). mTORC has been shown to be essential for Blimp-1 expression and plasma cell development^21,33^. To validate this downstream effect, we first confirmed the relationship between LARP4 and Blimp-1 expression in an in vitro differentiation system. Upon differentiation of B cells into plasma cells (by day 8), the expression of both LARP4 and Blimp-1 was significantly upregulated compared to undifferentiated B cells (Fig. 6a, b). On this basis, Western blot analysis revealed significantly reduced Blimp-1 protein levels in LARP4-deficient plasma cells differentiated at day 8 (Fig. 6c). Consistently, in the Bm12 lupus model and LPS+IL-4 in vitro experiments, flow cytometry showed that LARP4 deficiency led to a significant decrease in the frequency of Blimp-1^+^ plasma cells (Fig. 6d-g).

**Fig 6.**
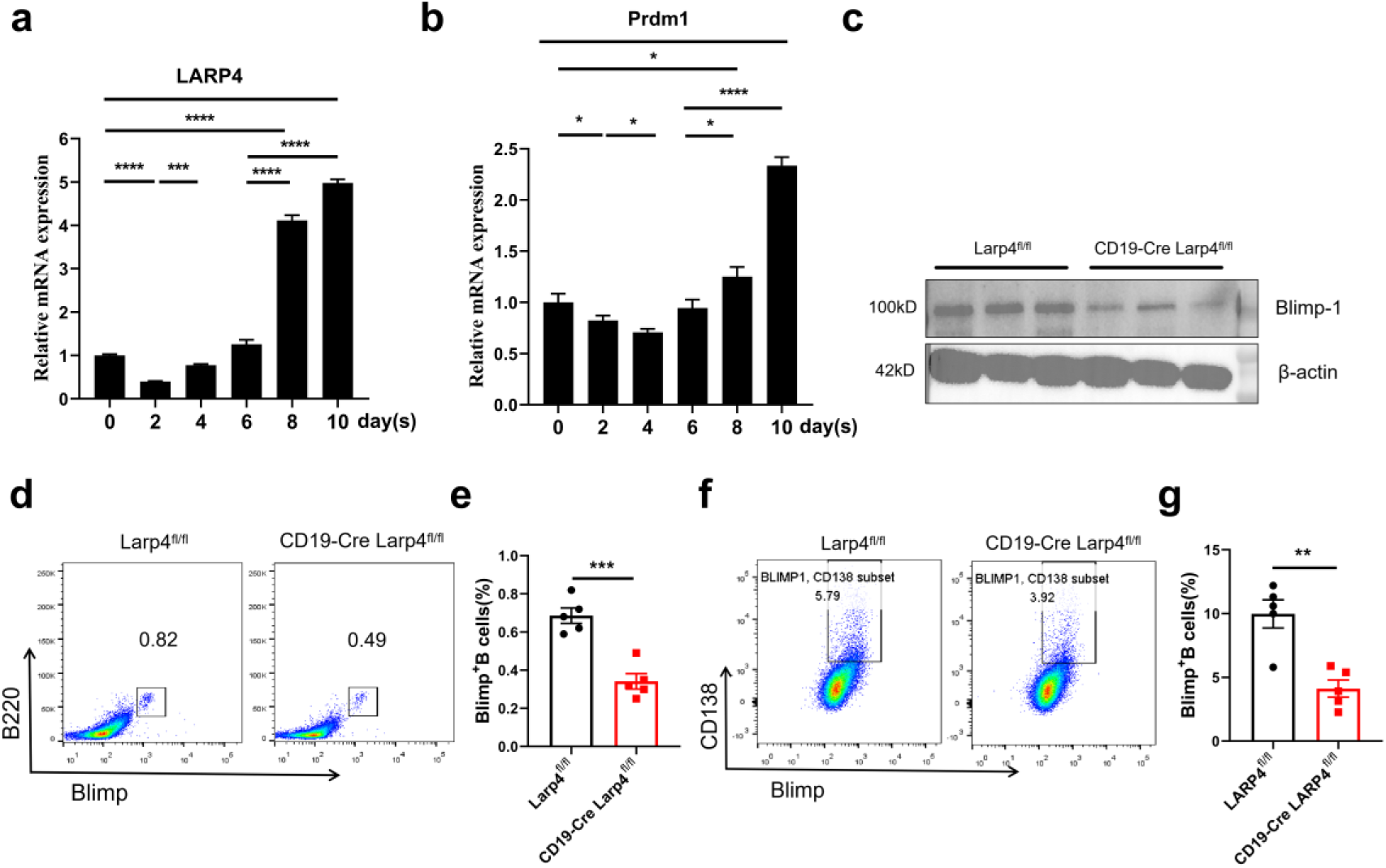
LARP4 deficiency suppresses Blimp-1 expression during plasma cell differentiation. **a, b.** RT-qPCR analysis of Larp4 **(a)** and Prdm1 **(b)** mRNA levels in pan-B cells stimulated with ImmunoCult™ Mouse B Cell Expansion Kit for 0,2, 4, 6, 8, and 10 days in vitro. **c.** Pan-B cells were stimulated for 10 days in vitro. The expression of Blimp-1 protein was presented by immunoblot (left) and its quantification (right), n = 3 biological replicates. **d,e.** Flow cytometry analysis of Blimp-1^+^ plasma cells in the spleens of Larp4^fl/fl^ and B-Larp4 CKO mice in the Bm12-induced SLE model (d) and the percentage of Blimp-1^+^ plasma cells is presented (e), n = 5 biological replicates. **f,g.** Flow cytometry analysis of Blimp-1^+^ plasma cells derived from Larp4^fl/fl^ and B-Larp4 CKO B cells stimulated with LPS and IL-4 for 3 days in vitro(f) and the percentage of Blimp-1^+^ plasma cells is presented (g), n = 4 biological replicates. Data are presented as mean ± SEM, ***p < 0.001, ****p < 0.0001 by two-tailed Student’s t-test.

In summary, these data establish a mechanistic cascade: LARP4 deficiency induces OXPHOS dysfunction and TCA cycle impairment, consequently impairing the de novo synthesis of PA. The subsequent reduction in PA levels culminates in decreased mTORC1 activity, which in turn attenuates the induction of Blimp-1, thereby offering a direct molecular rationale for the selective blockade of plasma cell differentiation observed in our study (Supplementary Fig. 8 Diagram of mechanism).

### 7. B-cell LARP4 deficiency selectively impairs IgG, but not IgM antibody responses

First, we quantified the frequencies of IgG1+ and IgM+ B cells in different models using flow cytometry. In the Bm12 lupus mouse model, the frequency of IgG1+ B cells in the spleen of B-LARP4 CKO mice was significantly reduced, while the frequency of IgM+ B cells did not differ significantly from that of the control group (Fig. 7a, b). This selective phenotype was consistently observed across the SRBC immunization mouse models (Fig. 7c, d), indicating that LARP4 deficiency specifically impairs the B cell population through class switch recombination (CSR), with minimal impact on unswitched IgM-producing B cells.

**Fig 7.**
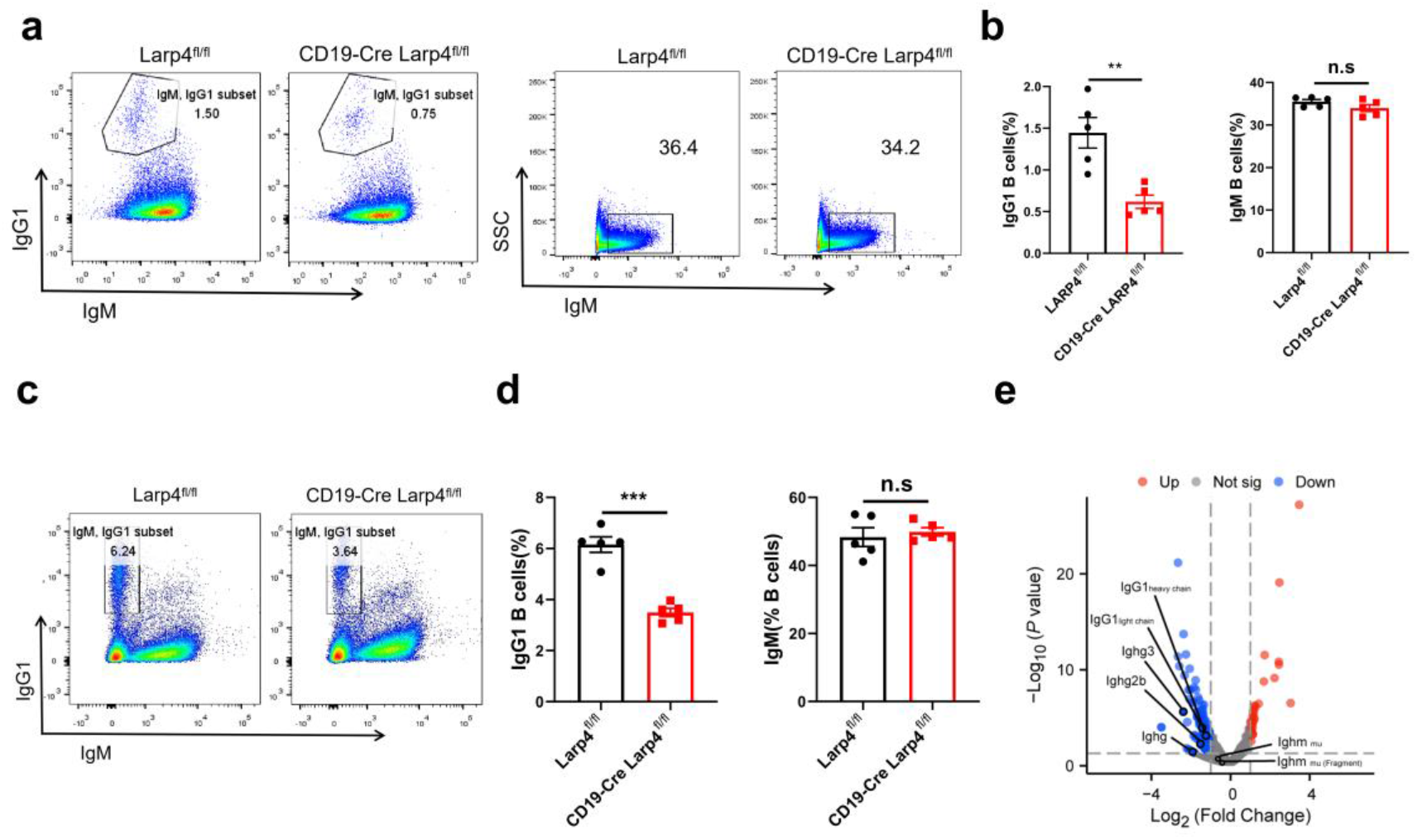
LARP4 deficiency selectively reduces IgG1⁺ but not IgM^+^ B cells. **a,b.** Flow cytometry analysis of IgG1⁺ and IgM^+^ B cells in the spleens of control Larp4^fl/fl^ and B-Larp4 CKO mice in the Bm12-induced SLE model **(a)** and the percentage is presented **(b)**, n =5 biological replicates. **c,d.** Flow cytometry analysis of IgG1⁺ and IgM^+^ B cells in the spleens of Larp4^fl/fl^ and B-Larp4 CKO mice upon immunization (i.p.) with SRBC at day 14 **(c)** and the percentage is presented **(d)**, n = 5 biological replicates. **e.** Proteomics analysis of IgG and IgM constant region proteins (IgG1, Ighg, Ighg2b, Ighg3, Ighm) in Larp4^fl/fl^ and B-Larp4 CKO plasma cells. Data are presented as mean ± SEM, *p < 0.05, **p < 0.01, ***p < 0.001by two-tailed Student’s t-test.

Subsequently, we measured serum antibody titers in immunized mice by ELISA. In both SRBC and NP-KLH T-cell-dependent antigen immunization models, the serum levels of IgG and IgG1 antibodies in B-LARP4 CKO mice were significantly lower than those in the control group. However, IgM antibody titers did not show a significant decrease and even exhibited a slight increase in some models (Fig. 8a-f). Consistently, in the supernatant of in vitro LPS+IL-4 cultures, the levels of IgG1 secreted by LARP4-deficient B cells were also significantly reduced, while IgM levels showed no significant change (Fig. 8g-i). These data are highly consistent with the flow cytometry findings, collectively demonstrating that LARP4 deficiency primarily affects the function of IgG-secreting plasma cells that have completed or are undergoing class switching.

**Fig 8.**
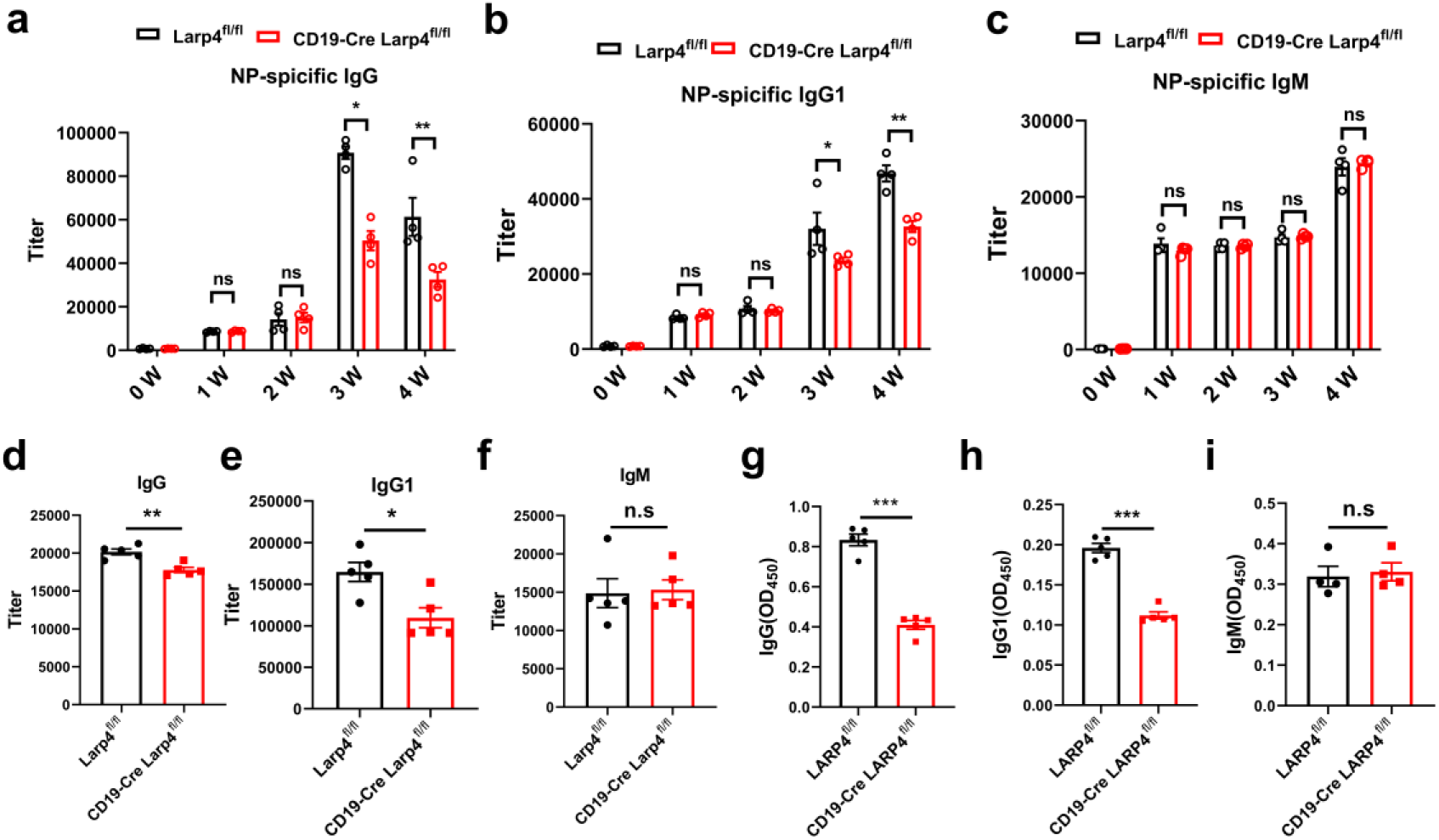
B cell-specific LARP4 deficiency selectively impairs IgG but not IgM antibody responses. **a-c.** ELISA analysis of serum IgG**(a)**,IgG1**(b)** and IgM**(c)** levels in Larp4^fl/fl^ and B-Larp4 CKO mice upon immunization (i.p.) with NP_33_-KLH, n =5 biological replicates. **d-f.** ELISA analysis of serum IgG**(d)**, IgG1**(e)** and IgM**(f)** levels in Larp4^fl/fl^ and B-Larp4 CKO mice upon immunization (i.p.) with SRBC at day 14, n =5 biological replicates. **g-i.** ELISA analysis of IgG**(g)**, IgG1**(h)** and IgM **(i)** levels in culture supernatants of Larp4^fl/fl^ and B-Larp4 CKO B cells stimulated with LPS and IL-4 for 3 days in vitro, n =4 biological replicates. Data are presented as mean ± SEM, *p < 0.05, **p < 0.01, ****p < 0.0001 by two-tailed Student’s t-test.

Ultimately, we substantiated the selective impact of LARP4 deficiency on IgG antibodies at the protein level through proteomic analysis. Specifically, the constant region proteins corresponding to IgG, including Ighg, IgG1, Ighg2b, and Ighg3, were significantly downregulated in LARP4-deficient plasma cells in the spleens of mice immunized to induce the Bm12 lupus model; conversely, the heavy chain protein Ighm corresponding to IgM exhibited negligible differences between the two groups (Fig. 7e). This result directly corroborates our previous flow cytometry and ELISA observations at the molecular level, providing strong evidence that LARP4 deficiency impairs the antibody synthesis capacity of B cells after CSR. Furthermore, non-targeted metabolomics analysis of LARP4-deficient plasma cells revealed that differential metabolites were significantly enriched in pathways such as the mTOR signaling pathway, branched-chain amino acid degradation, arginine and proline metabolism, and aminoacyl-tRNA biosynthesis (Fig. 5c). These findings suggest that metabolic alterations caused by LARP4 deficiency are intricately linked to CSR and its downstream antibody synthesis process^34,35^. In summary, these results systematically demonstrate, across cell frequency, antibody secretion function, and protein synthesis levels, that LARP4 deficiency selectively impairs IgG antibody production while preserving IgM responses, revealing the key role of LARP4 in B-cell class switching and IgG antibody generation.

### 8. LIPEP can mimic LARP4 deficiency and alleviate lupus nephritis in MRL/lpr mice

LARP4 shows co-localization with B cells and plasma cells in MRL/lpr lupus model mice. Multiplex immunofluorescence staining revealed that in the renal TLS of MRL/lpr mice, LARP4 expression primarily co-localized with B cells (B220⁺), whereas it showed no significant overlap with CD4^+^ and CD8^+^ T cells (Fig. 9a). Additionally, in the enlarged spleens, lymph nodes, and renal TLS of diseased MRL/lpr mice, LARP4 exhibited significant co-localization with plasma cells (CD138⁺) (Fig. 9b), suggesting that LARP4 expression is cell-type-specific.

**Fig 9.**
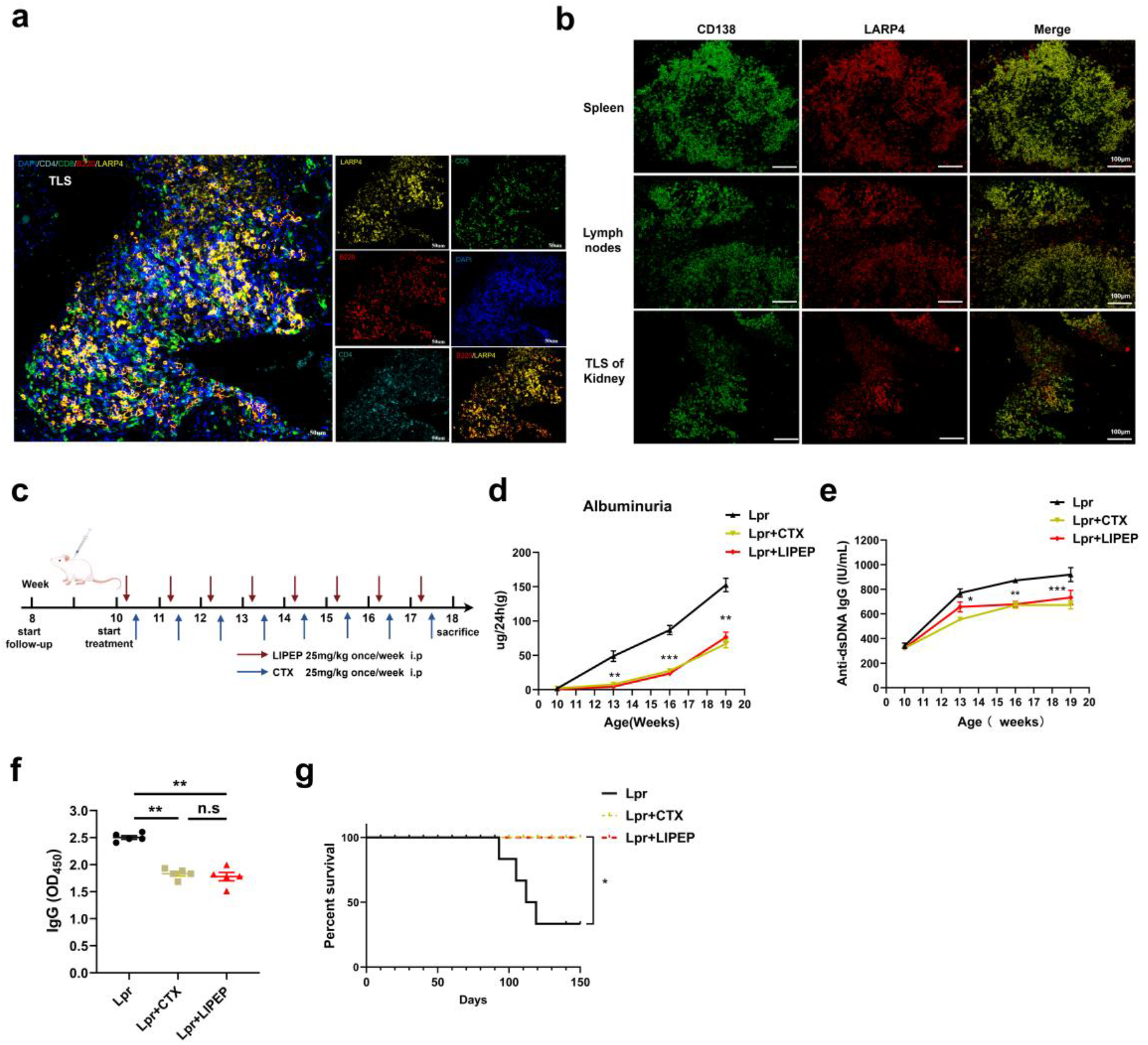
LARP4 colocalizes with B cells and plasma cells in MRL/lpr mice, and LIPEP treatment ameliorates systemic autoimmunity. **a.** Representative multiplex immunofluorescence staining of LARP4 (yellow), B220 (red), CD8(green), CD4(Cyan) and DAPI (blue) in kidney tertiary lymphoid structures (TLS) from MRL/lpr mice. Scale bar, 50 μm. **b.** Representative multiplex immunofluorescence staining of LARP4 (green), CD138 (red), and DAPI (blue) in the spleen, lymph node, and kidney TLS from MRL/lpr mice. Scale bar, 100 μm. **c.** Schematic of the experimental design. Female MRL/lpr mice (10-week-old) were randomly divided into three groups: disease control, CTX (25 mg/kg, i.p., once weekly), and LIPEP (25 mg/kg, i.p., once weekly). Treatment lasted for 8 weeks until 18 weeks of age. **d.** Total 24-hour Urine albumin was determined by ELISA. **e.** Serum levels of anti-dsDNA antibodies correlating with disease activity were measured by ELISA from week 10 to week 18. **f.** Serum levels of total IgG levels were detected by ELISA at 18 weeks of age. **g.** Kaplan-Meier survival curves of MRL/lpr mice in each group. n = 8–10 per group, log-rank test. *p < 0.05, **p < 0.01, ***p < 0.001, ****p < 0.0001 versus Lpr. Data are presented as mean ± SEM by one-way ANOVA with Tukey’s post-hoc test.

Based on these findings, we evaluated the therapeutic efficacy of a novel LARP4-inhibitory peptide, LIPEP, in the spontaneous lupus MRL/lpr mouse model, utilizing Cyclophosphamide (CTX) as a positive control. Ten-week-old female MRL/lpr mice were randomly assigned to three groups: the LIPEP treatment group, the CTX treatment group, and the disease control group. Treatment lasted for 8 weeks (intraperitoneal injection once a week, 25 mg/kg), during which various indicators were continuously monitored until 18 weeks of age (Fig. 9c). Compared to the disease control group, the LIPEP treatment group did not show significantly increased proteinuria during the treatment period (Fig. 9d). Furthermore, the degree of splenomegaly and lymph node enlargement, as well as splenocyte count, spleen weight, and spleen index, were significantly attenuated (Supplementary Fig. 9a-e). Serological tests showed that serum levels of anti-dsDNA antibodies and total IgG were significantly reduced in the LIPEP treatment group (Fig. 9e, f), while Complement C3 levels increased, and serum creatinine and blood urea nitrogen levels, reflecting renal function, also significantly decreased (Supplementary Fig. 9f-h). Additionally, the LIPEP treatment group demonstrated a significantly improved survival rate (Fig. 9g). In all the aforementioned renal function and systemic immune indicators, the therapeutic effects of LIPEP were comparable to those of CTX. These results indicate that LIPEP treatment ameliorated systemic autoimmune phenotypes and improved renal function.

Notably, LIPEP treatment alleviated renal histopathological damage and reduced immune complex deposition, demonstrating superior efficacy to CTX in glomerular deposits and interstitial inflammation. Histopathological evaluation using HE, PAS, and methenamine silver staining revealed that both LIPEP and CTX treatments significantly mitigated renal pathological damage (Fig. 10a). Compared to the control group, the LIPEP-treated group exhibited significantly reduced glomerulosclerosis, crescent formation, and interstitial fibrosis, at severity levels comparable to the CTX group (Fig. 10a). However, compared to the CTX-treated group, the LIPEP-treated group showed more pronounced reductions in scores for glomerular immune complex deposits and renal interstitial inflammatory cell infiltration (Fig.10a). Immunofluorescence staining further confirmed that both treatment groups exhibited a significant reduction in IgG and C3 deposits in the glomeruli compared to the control group, with the LIPEP-treated group demonstrating a marginally greater reduction than the CTX-treated group (Fig. 10b, c). LIPEP treatment also improved extrarenal manifestations, including dermatitis and vasculitis. Regarding the extrarenal pathological features characteristic of MRL/lpr mice, the severity scores for facial and back dermatitis and ear vasculitis in the LIPEP-treated group were significantly lower than those in the control group, with improvements surpassing those in the CTX-treated group (Fig. 10 d, e).

**Fig 10.**
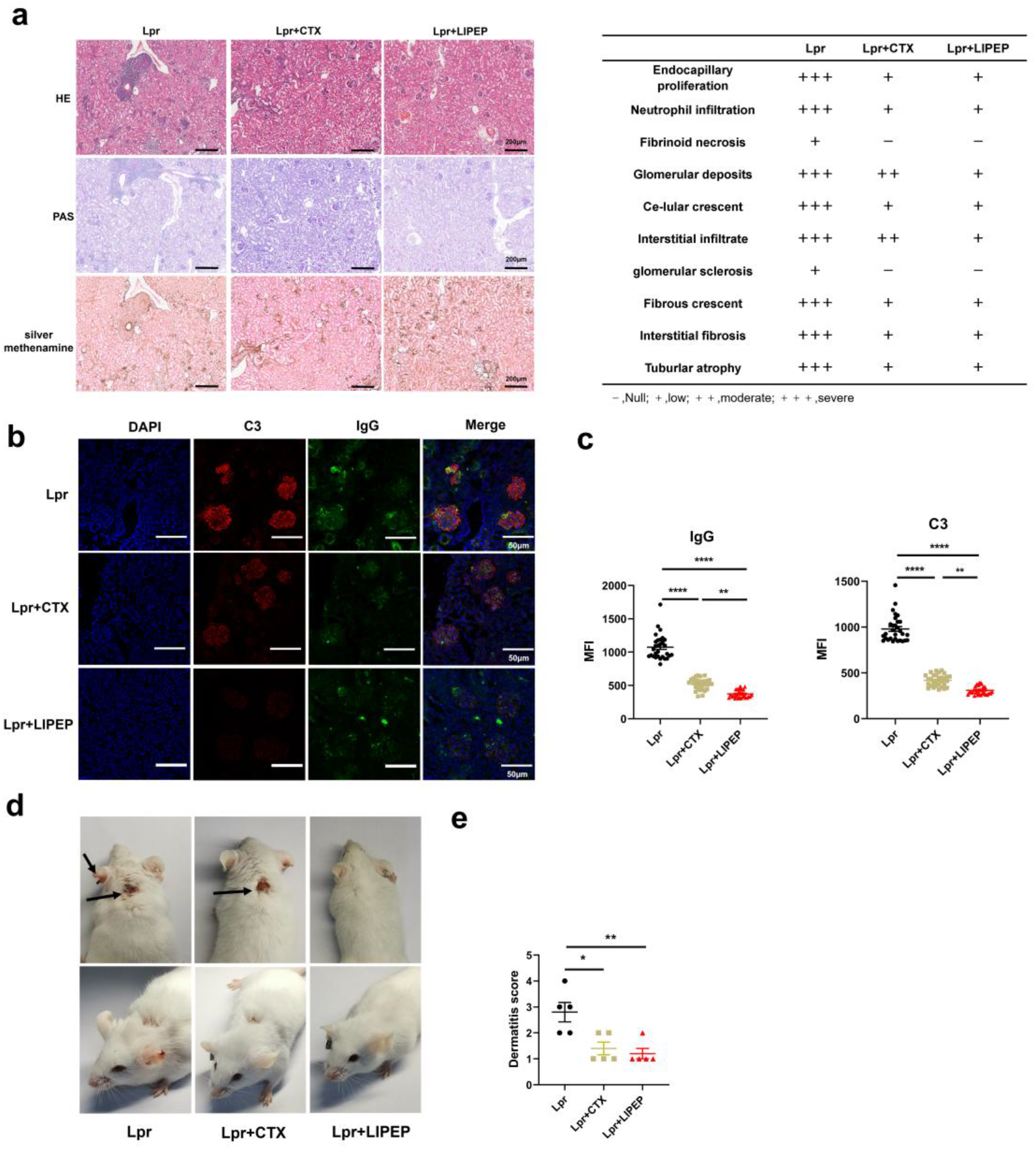
LIPEP treatment attenuates renal pathology and extrarenal manifestations in MRL/lpr mice. **a, b.** Representative H&E, PAS and silver methenamine-stained kidney sections **(a)** and histopathological scores **(b)** from DMSO, CTX, and LIPEP-treated MRL/lpr mice. Scores include glomerular sclerosis, crescent formation, interstitial fibrosis, glomerular deposits, and interstitial infiltrate, et al. **c,d.** Representative immunofluorescence staining of IgG **(c)** and C3 **(d)** deposition in kidney glomeruli (left) and quantification of fluorescence intensity (right). n = 6–8 per group. **e,f.** Representative photographs of facial and dorsal skin lesions and ear vasculitis **(e)** and severity scores of dermatitis and vasculitis **(f)** in DMSO, CTX, and LIPEP-treated MRL/lpr mice. n = 6–8, *p < 0.05, **p < 0.01, ***p < 0.01 versus Lpr; Data are presented as mean ± SEM by one-way ANOVA with Tukey’s post-hoc test.

In summary, the inhibitory peptide LIPEP, designed based on the expression characteristics of LARP4 in B cells and plasma cells, effectively recapitulates the therapeutic effects of LARP4 gene deletion in MRL/lpr lupus mice. LIPEP exhibits efficacy comparable to that of CTX in improving systemic immune responses, renal function, and kidney pathology, yet demonstrates superior efficacy in reducing glomerular deposits, inhibiting interstitial inflammation, and ameliorating extrarenal dermatitis and vasculitis.

## Discussion

Here, we report the key metabolic checkpoint function of the RNA-binding protein LARP4 in regulating the terminal differentiation of B cells into plasma cells. Using a B-cell-specific gene knockout mouse model, we found that LARP4 deficiency significantly attenuated Bm12-induced lupus nephritis without affecting germinal center B cell formation. We further demonstrated that LARP4 deficiency selectively inhibited plasma cell differentiation, and this effect was driven by mitochondrial dysfunction. Specifically, LARP4 deficiency resulted in impaired OXPHOS, which in turn compromised the synthesis of phosphatidic acid (PA). The reduction in PA directly suppressed mTORC1 activity, ultimately downregulating Blimp-1 expression and blocking the terminal differentiation of plasma cells. Mechanistically, we elucidated a complete upstream chain linking OXPHOS dysfunction to the mTORC1 signaling pathway through TCA cycle reprogramming and lipid metabolism dysregulation. From a translational perspective, we found that the LARP4-targeted inhibitory peptide LIPEP not only achieved efficacy comparable to cyclophosphamide in the MRL/lpr spontaneous lupus model but also demonstrated superior efficacy in reducing glomerular deposits, inhibiting interstitial inflammation, and improving extrarenal dermatitis. These findings establish LARP4 as a therapeutic target for B-cell-specific systemic lupus erythematosus and lay a theoretical foundation for developing more precise metabolic intervention strategies for autoimmune diseases.

This study revealed via transcriptome analysis that LARP4 is specifically highly expressed in B cells of SLE patients and correlates positively with disease activity (SLEDAI score). Although the function of LARP4 in T cells has been reported, including its involvement in activation, differentiation, and exhaustion processes ^27,30^, our data suggest that its B-cell-specific expression may have unique pathogenic significance in lupus pathogenesis. To investigate this hypothesis, we constructed conditional knockout mice targeting specific cell lineages. In the Pristane-induced lupus model, T-cell-specific LARP4 deletion had no significant effect on disease progression. In contrast, B-cell-specific LARP4 deletion significantly alleviated multiple phenotypes of lupus nephritis, including splenomegaly, anti-dsDNA antibody production, renal immune complex deposition, and glomerular injury. This result localizes the pathogenic role of LARP4 to B cells, rather than T cells, suggesting it may serve as a B-cell-selective therapeutic target.

At the cellular level, we found that LARP4 deletion significantly inhibited B cell differentiation into plasma cells but did not affect the number, size, or proliferation of germinal center (GC) B cells. This stage-specific phenotype contrasts with previously reported B-cell intrinsic defects. For example, the deletion of MCT1 impairs both GC B cell and plasma cell differentiation^18^, while the deletion of EZH2, although also preferentially blocking plasma cell generation, exerts a more nuanced regulation on GC B cells^36^. These differences may arise from the distinct metabolic pathways regulated by different molecules. One possible explanation is that GC B cells primarily rely on fatty acid oxidation (FAO) to drive mitochondrial respiration for their energy needs^37,38^, utilizing glucose and amino acids for biosynthesis—a metabolic pattern that may not critically depend on LARP4. In contrast, plasma cells, which require massive synthesis and secretion of antibodies, experience a sharp increase in energy demand and are highly dependent on oxidative phosphorylation (OXPHOS)^39,40^. Therefore, the OXPHOS deficiency resulting from LARP4 deletion is exacerbated during the plasma cell differentiation stage—a process heavily reliant on OXPHOS—ultimately acting as a limiting factor for terminal differentiation. This finding aligns with the unique metabolic characteristics of plasma cells and partially explains why certain metabolism-related defects exhibit stage-selective effects.

A key finding of this study lies in elucidating how OXPHOS dysfunction affects downstream signaling through metabolic reprogramming. Metabonomics data indicate that in LARP4-deficient plasma cells, the TCA cycle exhibits characteristic changes: levels of metabolites upstream of succinate dehydrogenase (SDH), such as citrate, are reduced, while downstream products, such as fumarate and malate, accumulate. This pattern has been reported in models of B-cell mitochondrial dysfunction^12,41^, consistent with the global downregulation of OXPHOS subunits observed in our study. This TCA cycle imbalance further affects lipid metabolism, leading to an overall reduction in phosphatidic acid (PA) levels. PA is not only a key component of cell membranes but also a direct activator of the mTORC1 complex. Our data show that a decrease in saturated and monounsaturated short-chain PA, accompanied by an accumulation of long-chain polyunsaturated PA, indicates a dysregulation in fatty acid metabolism. These findings provide a direct upstream metabolic mechanism underlying the reduced mTORC1 activity induced by LARP4 deficiency. Interestingly, this connection between metabolism and signaling is cell-type specific. In T cells, LARP4 deficiency similarly downregulates OXPHOS but reportedly reduces ROS levels^28^, a finding that contrasts with the metabolic stress features observed in our B-cell model of OXPHOS dysfunction. This difference may reflect fundamental distinctions in the primary functions and energy demands of the two cell types, also highlighting the necessity of metabolic interventions tailored to different immune cells.

At the effector function level, we observed that LARP4 deficiency selectively impairs the production of IgG, but not IgM antibodies. The production of IgG antibodies depends on T cell-dependent antigen responses and class switch recombination, a process that requires plasma cells to provide sufficient energy and maintain mTORC1 activity. Our previous results demonstrated that LARP4 deficiency inhibits the secretory function of IgG plasma cells by reducing PA levels, decreasing mTORC1 activity, and downregulating Blimp-1 expression, whereas IgM responses remain largely unaffected. From a clinical translation perspective, selectively reducing pathogenic IgG autoantibodies (such as anti-dsDNA) while preserving natural IgM, which plays critical roles in immune regulation and apoptotic cell clearance, is an ideal therapeutic strategy. Although traditional broad-spectrum immunosuppressants such as cyclophosphamide (CTX) can effectively suppress IgG, they also inhibit IgM and its associated protective functions. Our LIPEP therapeutic peptide mimics the efficacy of gene knockout in the MRL/lpr model: it not only achieves effects comparable to CTX in protecting renal function and prolonging survival but also demonstrates superior efficacy in reducing glomerular deposition, inhibiting interstitial inflammation, and improving extrarenal dermatitis and vasculitis. This suggests that LIPEP reduces the non-specific immunosuppression caused by CTX by more specifically targeting B cells or plasma cells, thereby offering enhanced selectivity.

In summary, our work integrates LARP4, mitochondrial dysfunction, PA metabolism, mTORC1 signaling, and antibody production at the molecular, metabolic, and functional levels, highlighting LARP4’s potential as a B cell-specific therapeutic target. As a lead compound targeting this pathway, LIPEP yields preliminary data supporting its efficacy and selectivity for the development of safer immunotherapies for SLE.

## Methods

### Mice

For conditional knockouts, loxP sites are typically inserted into introns downstream of the exon containing the ATG start codon, where the excision of the flanked exon(s) induces a protein reading frame shift. After reviewing the gene structure and exon sizes, we identified exons 3(ENSMUSE00000344606) and 4 (ENSMUSE00000489242) as suitable candidates for conditional excision; the deletion of exons 3 and 4 would result in a 60 aa (51 native aa plus 9 in-frame aa) truncated protein potentially subject to nonsense-mediated decay (NMD). This strategy enables the generation of LARP4-/-mice (KO) via CRISPR/Cas9-mediated total knockout or Cre-loxP-mediated conditional excision of exons 3 and 4. Given the substantial size of introns 2 and 4, the insertion of loxp sites is unlikely to interfere with mRNA splicing. To minimize the risk of disrupting Larp4 expression, both loxP sites were inserted into non-conserved regions. Larp4^fl/fl^ mice were provided by BIOCYTOGEN Technology Ltd. (Beijing, China) and were backcrossed to C57BL/6J mice for ten generations. Larp4-floxed mice were crossed with Cd4-Cre mice (BIOCYTOGEN Technology Ltd, Beijing, China) and Cd19-Cre mice (Shanghai Southern Model Biotechnology Ltd, Shanghai, China) to generate CD4-Larp4 CKO and B-Larp4 CKO mice. MRL-lpr female mice and BM12 (JAX1162) mice were purchased from the Jackson laboratory. All mice were housed in a specific pathogen-free (SPF) environment under a 12-hour light/dark cycle in a temperature-controlled room with ad libitum access to food and water. All mice were maintained under specific pathogen-free conditions in the animal facility of Army Medical University and handled in accordance with government and institutional animal welfare guidelines. All animal experiments were approved by the Institutional Animal Ethics Committee.

### BM12-induced lupus-like mouse model

To induce lupus-like model, splenocytes (1 x 10^7^ per mouse) from age and gender-matched BM12 mice were adoptively transferred to 6-8-week wild-type C57BL/6 mice for two times on a weekly basis. Mice were sacrificed at the week 4 for the assessment of autoantibodies, glomerular immune complex deposition, and renal histopathological damage.

### Non-targeted metabolomics analysis of mouse plasma cells

Comprehensive metabolomic analysis of plasma cells from Larp4fl/fl and CD19-Cre Larp4fl/fl mouse groups in mice. The specific steps for sample preparation were as follows: 50 mg of sample was weighed, and 1000 μL of an extraction solution consisting of methanol, acetonitrile, and water (2:2:1) was added, followed by vortexing for 30 seconds; then the sample was homogenized using a grinder for 10 minutes and sonicated for 10 minutes in an ice water bath; subsequently, the sample was centrifuged at 4°C and 12,000 rpm for 15 minutes; finally, the supernatant was collected and subjected to LC-MS/MS analysis.After normalizing the original peak area information with the total peak area, the follow-up analysis was performed. Principal component analysis and Spearman correlation analysis were used to judge the repeatability of the samples within group and the quanlity control samples. The identified compounds are searched for classification and pathway information in KEGG, HMDB and lipidmaps databases.According to the grouping information, calculate and compare the difference multiples, T test was used to calculate the difference significance pvalue of each compound. The R language package ropls was used to perform OPLS-DA modeling, and 200 times permutation tests was performed to verify the reliability of the model. The VIP value of the model was calculated using multiple cross-validation. The method of combining the difference multiple, the P value and the VIP value of the OPLS-DA model was adopted to screen the differential metabolites. The screening criteria are FC>1, P value<0.05 and VIP>1. The difference metabolites of KEGG pathway enrichment significance were calculated using hypergeometric distribution test.

### Transcriptomic and Proteomic Analysis

RNA sequencing (RNA-seq): Total RNA was extracted from sorted plasma cells, and libraries were constructed using the Illumina TruSeq mRNA Library Prep Kit, followed by 150 bp paired-end sequencing on the Illumina NovaSeq 6000 platform. Sequencing data were aligned to the mouse reference genome, and differentially expressed genes were screened with criteria of |log2FC| ≥ 1 and FDR < 0.05. Gene Ontology (GO) enrichment analysis was performed to identify enriched pathways. For proteomic analysis, sorted plasma cells were lysed in RIPA buffer, then reduced, alkylated, and digested with trypsin, followed by LC-MS/MS analysis using TMT labeling or label-free quantification methods. Data was searched, identified, and quantified using MaxQuant software, aligned to the UniProt mouse protein database.

### Flow cytometry and cell sorting

Single-cell suspensions were prepared from the spleen and bone marrow. Following Fc receptor blocking with anti-CD16/32, surface staining was performed according to standard protocols. Data acquisition was conducted using the BD LSRFortessa system and analyzed with FlowJo software (v10.8, TreeStar). Relevant antibodies included anti-B220 , anti-CD19 , anti-CD138, anti-GL-7, anti-CD95, anti-CD4, anti-CXCR5, anti-IgG1, and anti-IgM (all purchased from BD Biosciences or BioLegend). For intracellular staining, cells were fixed with a fixation/permeabilization solution (BD Biosciences) at 4°C for 1 hour, and antibodies included anti-Blimp-1 and anti-pS6 (S235/236) (BioLegend). For transcriptomic, metabolomic, and proteomics analyses, splenic B220⁻CD138⁺ plasma cells were sorted on a BD FACSAria III sorter with a purity greater than 95%.

### Flow cytometry analyses mitochondria mass, ROS and transmembrane potential

Mitochondrial reactive oxygen species (ROS) levels were detected using the MitoSOX Green mitochondrial superoxide indicator (Thermo Fisher Scientific, M36005), mitochondrial membrane potential was assessed with the MitoTracker Red CMXROS (Thermo Fisher Scientific, M46752), and mitochondrial mass was measured using the MitoTracker Green FM (Thermo Fisher Scientific, M46750). Co-staining with these two dyes enabled the evaluation of membrane potential changes per unit mitochondrial mass. Single-cell suspensions were incubated with these dyes at 37°C in a 5% CO2 incubator for 30 minutes, washed, resuspended in flow cytometry buffer, and subjected to flow cytometric analysis.

### Mouse immunization and ELISA assay

For TD immunization, 6-week-old mice were intraperitoneally injected with 100 mg of NP^33^-KLH emulsified in Alum adjuvant. Mice were sacrificed 4 weeks after immunization. Blood was collected weekly post-immunization, and serum was harvested and stored frozen until analysis. To detect NP-specific or anti-dsDNA antibodies, 4 µg/mL dsDNA or 5 mg/ml NP-BSA in PBS was coated onto 96-well ELISA, plates (MCE, HY-NP141) overnight and washed. Affinity-purified goat anti-mouse IgM and goat anti-mouse IgG-Fc fragment antibodies were used for coating to detect naturally occurring IgM and IgG in serum. Each serum sample was serially diluted starting from 1:500, added to the plate, and incubated at room temperature for 2 hours. After PBST washing, the plate was further incubated with HRP conjugated goat anti-mouse IgM,IgG,IgG1 at 1:6,000 for 1h before development with TMB Single-Component Substrate solution (423001,BioLegend) .For PC examination,, 6-week-old mice were intraperitoneally injected with sheep red blood cells (S9702-100 ml,Solarbio), Mice were analyzed on day 14 after immunization.

### Immunohistochemistry for immune complexes in glomeruli

Kidneys were obtained from BM12-induced lupus,Pristane and MRL/Lpr mice,fixed with 4% paraformaldehyde, cryoprotected with 30% sucrose solution and frozen in O.C.T compound at −80 ℃ , and then sliced into 5μm sections for immunohistochemistry. Kidney slides were rehydrated and blocked, then incubated with rabbit anti-mouse C3-specific antibody at 4°C overnight. Stained with fluorescence-conjugated goat anti-rabbit IgG for 2 h.stained with fluorophore-coupled goat anti-mouse IgM and IgG specific antibodies for 12h.Sections were imaged with the ultra-high-resolution laser confocal microscope (ZISS 800).

### Western blotting

Cell samples were lysed using RIPA buffer (containing protease inhibitor cocktail) on ice for 30 min, centrifuged at 12,000 g for 15 min at 4 ℃, and the supernatant was collected. Protein concentration was determined using the BCA method. Equal amounts of protein were separated by 10% SDS-PAGE and wet-transferred to a PVDF membrane. The membrane was blocked with 5% non-fat milk at room temperature for 1 h, then incubated overnight at 4 ℃ with the following primary antibodies: rabbit anti-LARP4 (1:3000, Novus Biologicals NBP2-22254), rabbit anti-Blimp-1 (1:1000, CST 9115T), and rabbit anti-β-actin, which served as an internal control. After washing, the membrane was incubated with HRP-conjugated goat anti-rabbit secondary antibody at room temperature for 1 h, visualized using enhanced chemiluminescence (ECL) substrate, and images were captured with a gel imaging system.

### Multiplex immunofluorescence staining

Paraffin-embedded kidney, spleen, or lymph node tissue sections were routinely deparaffinized and rehydrated through a graded ethanol series, followed by high-pressure heat-induced antigen retrieval in citrate buffer at pH 6.0/9.0. After cooling, the sections were immersed in 3% H₂O₂ for 15 min to block endogenous peroxidase and were subsequently blocked with PBS containing 10% goat serum at room temperature for 30 min. Subsequently, multiple rounds of TSA staining were performed. Sections were first incubated with rabbit anti-LARP4 primary antibody (1:600, Novus NBP1-80889) overnight at 4°C. After washing, they were incubated with HRP-conjugated goat anti-rabbit secondary antibody at room temperature for 30 min, followed by reaction with TSA fluorescent dye at room temperature for 10 min, during which peroxidase covalently deposited fluorescently labeled tyramide at the antigen site. After one round was completed, the sections underwent microwave treatment in pH 6.0 citrate buffer at 95°C for 15 min to remove the antibody-dye complex. The following primary antibodies were then used sequentially for staining: rabbit anti-CD138 (1:2000, Proteintech 10593-1-AP), rabbit anti-B220 (1:200, Novus NBP2-53303), rabbit anti-CD4 (1:200, CST 25229), and rabbit anti-CD8 (1:400, CST 98941), each assigned to a different TSA fluorescence channel. After all staining was completed, the sections were counterstained with DAPI and mounted with an anti-fade mounting medium. A multispectral imaging system was used to scan and capture multi-channel images to evaluate the co-localization of each labeled molecule.

### H&E and PAS taining

Kidney tissues were fixed with 4% paraformaldehyde, embedded in paraffin, and sectioned (3 μm) for H&E and PAS staining.The glomerular injury score was semi-quantitatively assessed based on glomerular endothelial cell proliferation, crescent formation, inflammatory cell infiltration, and renal tubulointerstitial injury.

### Renal function analysis

24-hour urine was collected from mice, and urinary protein levels in 4-hour urine samples were quantified using an automatic biochemical analyzer; urinary albumin concentrations were assessed using a commercial ELISA kit (Elabscience). Serum was collected from mice, and levels of blood urea nitrogen, serum creatinine, and complement C3 were determined using specific assay kits according to the manufacturers’ instructions.

### LIPEP treatment in MRL/lpr mice

Female MRL/lpr mice (10 weeks old) were randomly assigned to three groups (n = 8–10 per group): (1) LIPEP treatment group, (2) cyclophosphamide (CTX) positive control group, and (3) disease control group (DMSO). LIPEP (25 mg/kg body weight, prepared in 2% mannitol) was administered via intraperitoneal (i.p.) injection once per week for 8 consecutive weeks (from week 10 to week 18 of age). CTX was administered at a clinically relevant dose of 25 mg/kg body weight via i.p. injection once per week over the same treatment period.

During the treatment period, body weight, proteinuria, and survival were assessed weekly. At the end of the study (week 18), mice were euthanized. Blood was collected for serum separation, and sera were stored at –80°C for subsequent antibody detection. Spleens and lymph nodes were harvested for cellular analysis, and kidneys were collected for histopathological assessment and immunofluorescence staining. Skin lesions (dermatitis) and ear vasculitis were scored macroscopically.

### Public database analysis

The GSE10325 dataset comprises RNA-seq data from PBMCs of healthy controls and SLE patients. The expression of LAPR4 was retrieved from B cells, CD4^+^T cells, and myeloid cells. Statistical analyses were performed in R using the Wilcoxon signed-rank test to compare the two groups, with P<0.05 considered statistically significant.

### Statistical analysis

Statistical analysis was performed using GraphPad Prism 11. Comparisons between two groups were conducted using unpaired t-test or Mann-Whitney U test. Comparisons among multiple groups were performed using one-way analysis of variance (ANOVA).Data are presented as mean±SD, and statistical significance is denoted with * (P < 0.05), ** (P < 0.01), and ***(P < 0.001) in the figures. Experiments were repeated two to three times.

## Data availability

All relevant data from this study are available from the corresponding authors upon reasonable request.

## Funding

This work was supported by grants from the Major Research Plan of the National Natural Science Foundation of China (No. 92569203 and 92269110), the Special Program of the National Natural Science Foundation of China (No. 32141005), the General Program of the National Natural Science Foundation of China (No. 32170887 and 82271846), as well as the Chongqing Talent Program of China (cstc2022ycjh-bgzxm0020), Chongqing Natural Science Foundation Key Project(NO.CSTB2024NSCQ-KJFZZDX0049), Chongqing Joint Medical Research Project of Municipal Health Commission and Municipal Science and Technology Bureau(NO.2026GDRC006), Special fund for performance incentive guidance of research institutions in Chongqing (cstc2021jxjl130036) and Chongqing International Institute for Immunology Project (2022YJC01). The funders had no role in the study design, data analyses, or the decision to publish.

## Competing Interests Statement

All the authors have no competing interests.

---

**Supplementary Fig 1.**
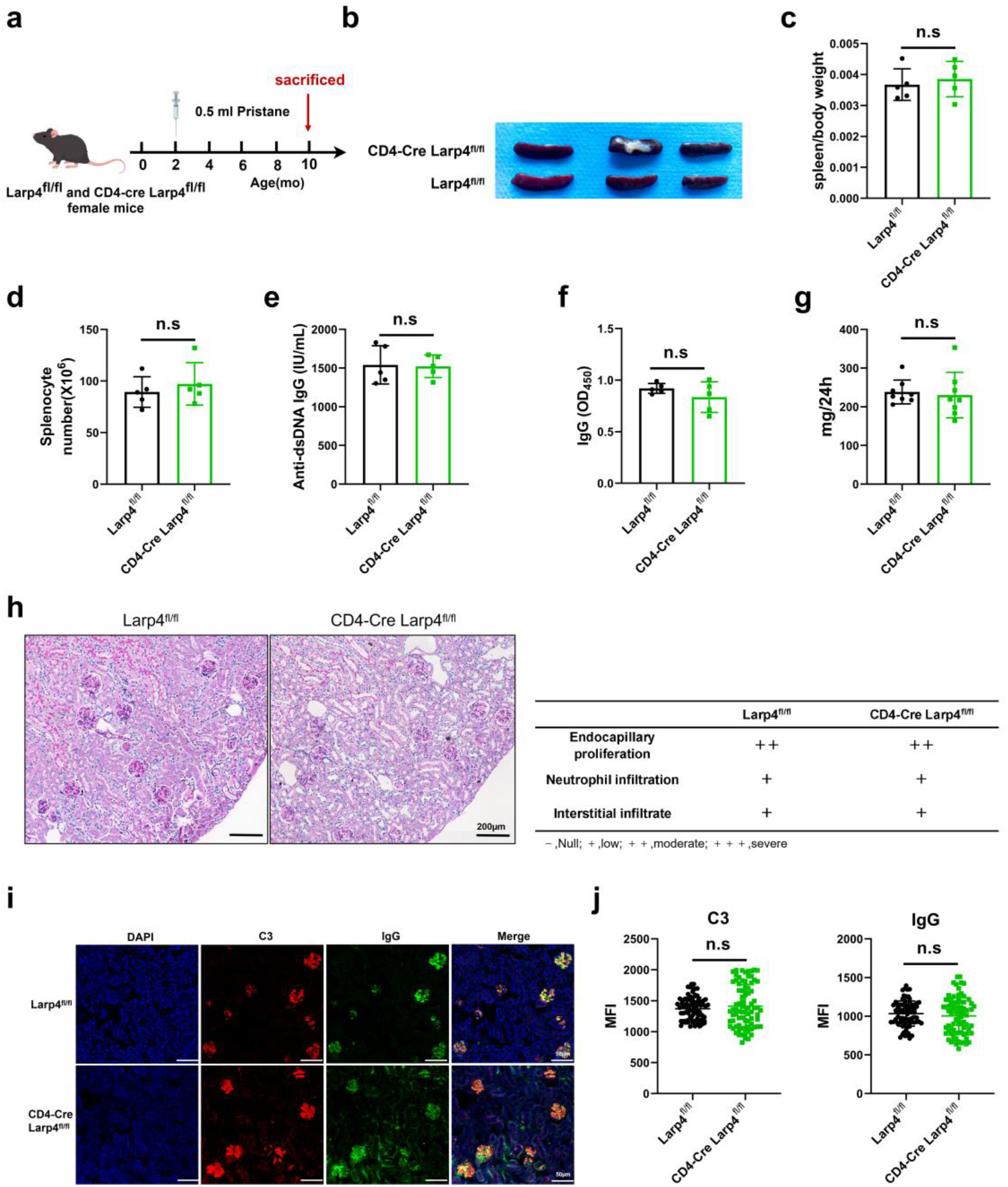
CD4^+^T cells-specific LARP4 deficiency does not protect against pristane-induced lupus nephritis. **a.** Schematic of the experimental design for establishing the pristane-induced lupus model in CD4-Larp4 CKO (Larp4^fl/fl^CD4-Cre) and Larp4^fl/fl^ mice, n = 6 biological replicates. **b-d.** Spleen size **(b)**, spleen index (spleen weight/body weight, **c**), and total splenocyte counts **(d)** in Larp4^fl/fl^ and CD4-Larp4 CKO mice at 6 months post-pristane injection. **e,f.** Serum anti-dsDNA IgG and total IgG levels were measured by ELISA.**g.** 24-hour urinary protein levels were measured by Urine protein quantification kit. **h.** Representative PAS-stained kidney sections (left, scale bar=200μm) and glomerular histopathology scores (right), including glomerular hypercellularity, inflammatory infiltration, and tubular interstitial injury. **i, j.** Representative immunofluorescence staining of IgG and C3 deposition in kidney glomeruli (**i,** scale bar = 200 μm) and quantification of fluorescence intensity **(j)**. Data are presented as mean ± SEM. ns, not significant by two-tailed Student’s t-test or Mann-Whitney U test.

**Supplementary Fig 2.**
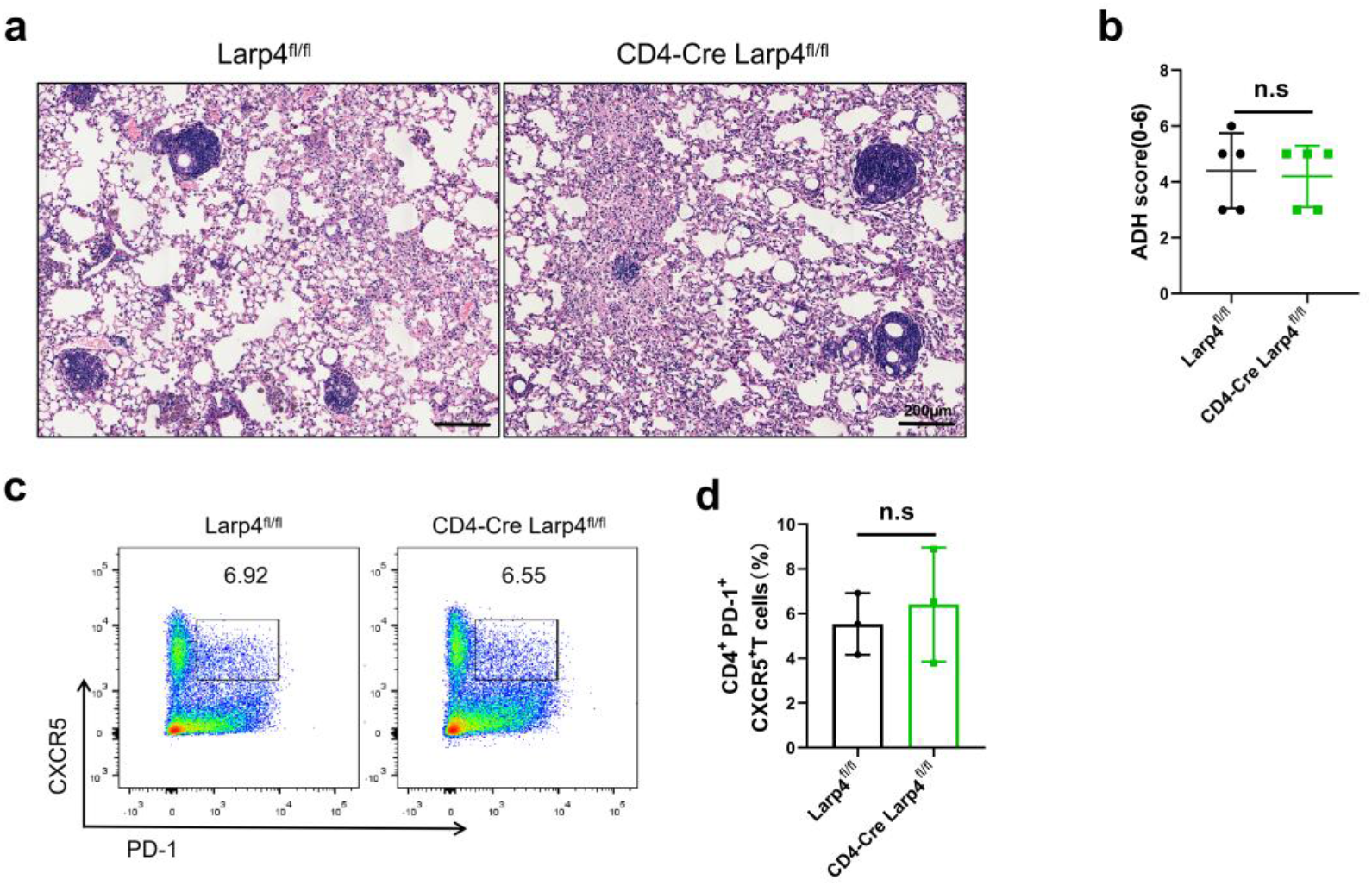
CD4^+^T cell-specific LARP4 deletion does not affect pulmonary pathology in pristane-induced lupus mice. **a,b.** Representative H&E-stained lung sections from Larp4^fl/fl^ and CD4-Larp4 CKO mice at 6 months post-pristane injection (**a,** scale bar = 200 μm). Quantification of ADH severity score **(b)**. **c,d.** Representative flow cytometry plots **(c)** and quantification **(d)** of CD4⁺T cell subsets Tfh in the spleens of pristane-treated mice. Data are presented as mean ± SEM. ns, not significant by two-tailed Student’s t-test.

**Supplementary Fig.3.**
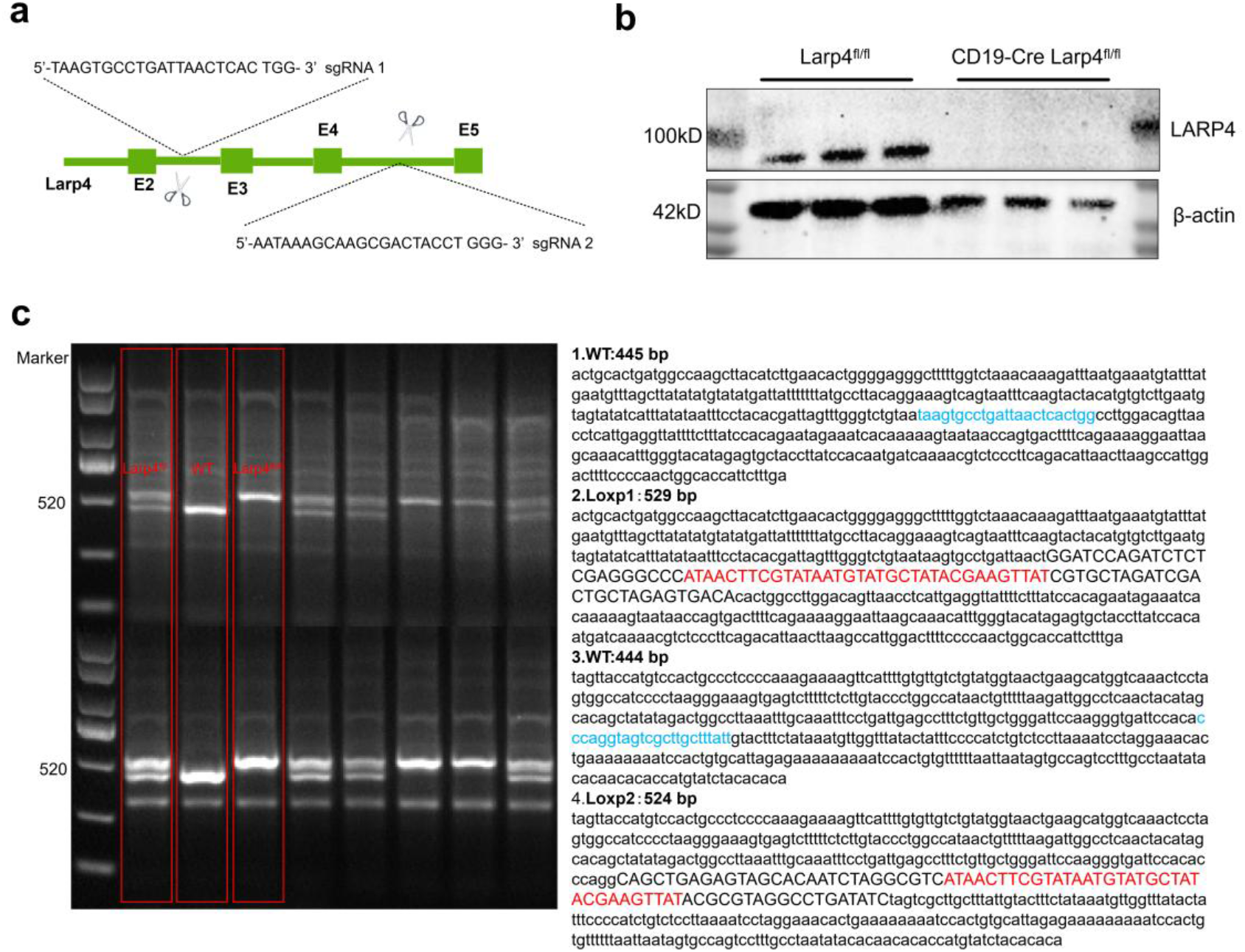
The Larp4 mice were obtained with CRISPR-Cas9 technology. **a.** The schematic diagram of the sgRNA sites at Larp4 allele. **b.** Western blotting (WB) showed the expression of the LARP4 protein in B cells from CD19-Larp4 CKO and Larp4^fl/fl^ mice. **c.** PCR identification analysis of targeted alleles. Expected fragment size: 1-WT = 445 bp, 2-LoxP 1 = 529 bp, 3-WT = 444 bp, and 4-LoxP 2 = 524 bp.

**Supplementary Fig 4.**
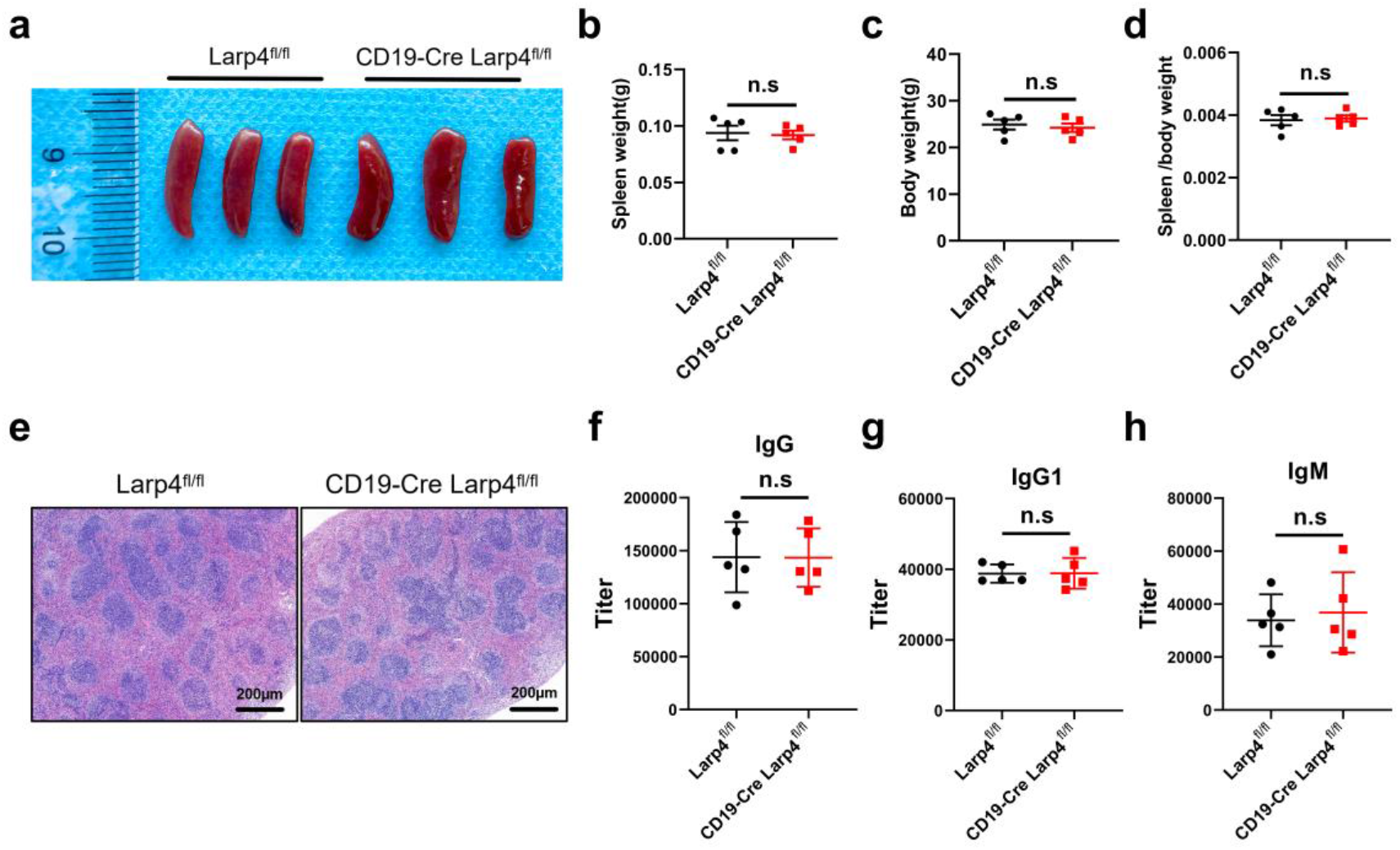
LARP4 deficiency does not affect spleen morphology, weight or the serum innate antibodies homeostasis. **a-e.** In non-immunized Larp4^fl/fl^ and CD19-cre Larp4^fl/fl^ mice, spleen size **(a)**, spleen weight **(b)**, body weight **(c)**, and the spleen-to-body weight ratio **(d)** was presented (n = 5 biological replicates per group). The morphology of the red and white pulp of the spleen was analyzed by H&E. Scale Bar, 200μm **(e)**. Titer of innate antibody IgM, IgG or IgG1 isotype were detected by ELISA, n = 3 biological replicates**(f)**. Data (a, b, c, d, f) are representative one of three independent experiments. Data are presented as mean ± SEM of 5-6 mice, unpaired two-tailed t-test, n.s, not significant.

**Supplementary Fig 5.**
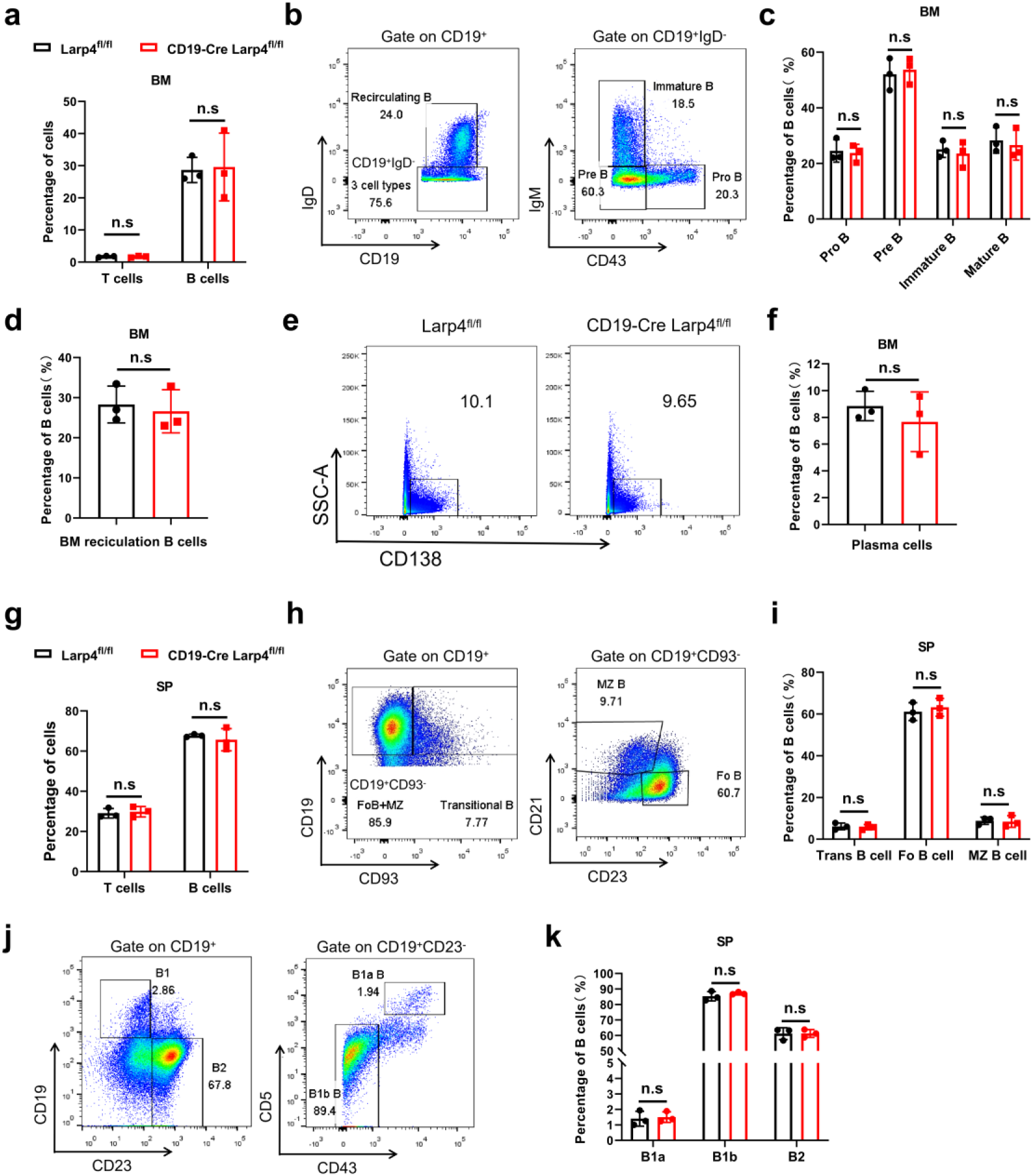
LARP4 deficiency does not impair the B cells development. **a-d.** Flow cytometry analysis of the frequencies of T cells, B cells, immature B cells, pro-B, pre-B, recirculating mature B cells, follicular B cells, marginal B cells, transitional B cells, B1a, B1b and B2 cells , Plasma cells in bone marrow (BM) **(a-f)** or spleen (SP) **(g-k)**, respectively, n = 3 biological replicates. Data are presented as mean±SEM of 3 mice, unpaired two-tailed t-test, n.s, not significant.

**Supplementary Fig 6.**
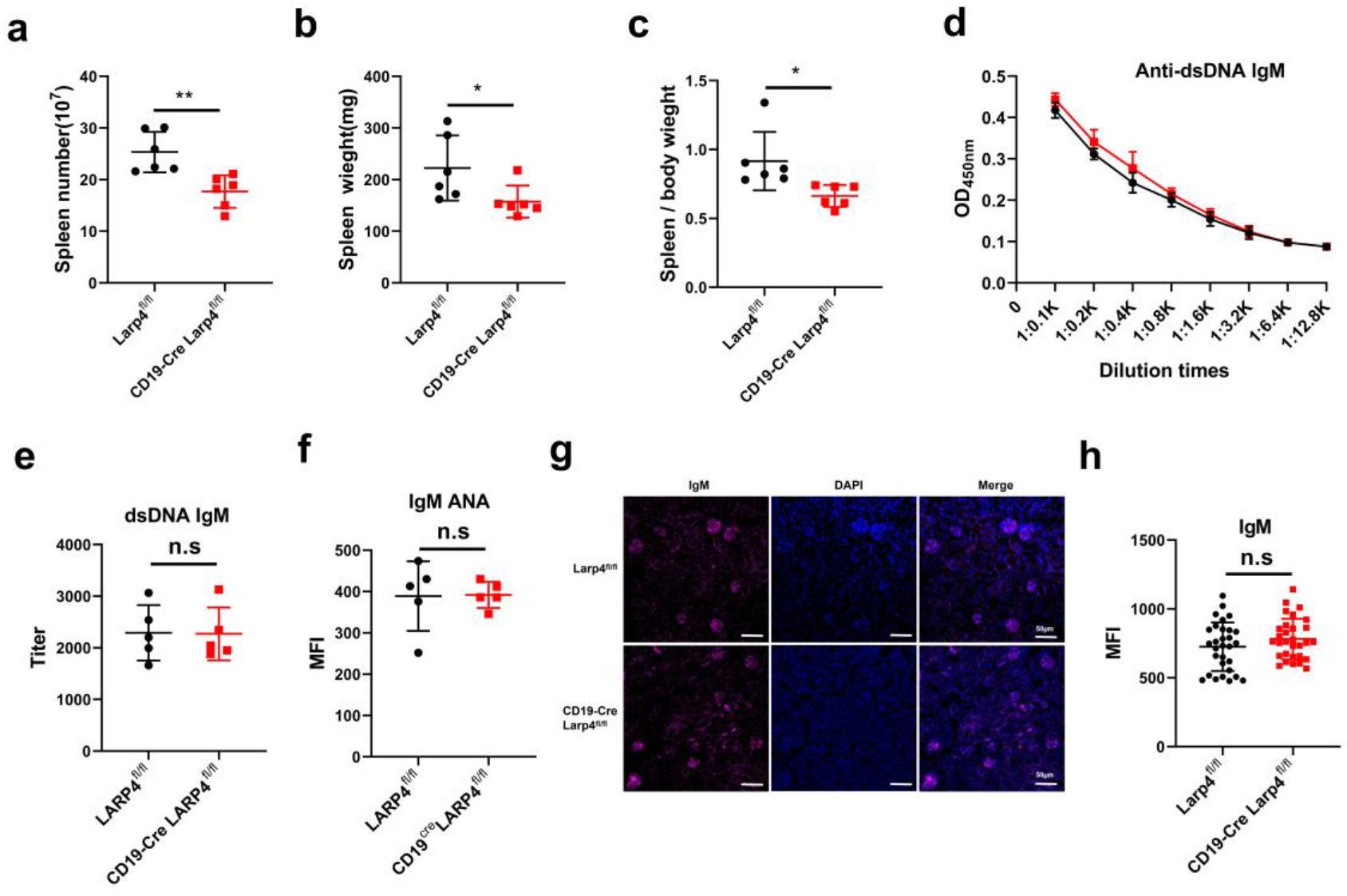
B cell-specific LARP4 deficiency reduces splenomegaly and IgG autoantibodies while sparing IgM responses. **a-c.** Spleen weight **(a)**, splenocyte counts **(b)**, and spleen index **(c)** in Larp4^fl/fl^ and B-Larp4 CKO mice. **d,e.** OD450 nm of serum anti-dsDNA IgM and their titers measured by ELISA. **f.** Serum levels of anti-ANA IgM measured by Flow cytometry. **g,h.** Representative immunofluorescence staining of IgM deposition in kidney glomeruli **(g)** and quantification of fluorescence intensity **(h)**. Data are presented as mean ± SEM. ns, not significant; *p < 0.05, **p < 0.01 by two-tailed Student’s t-test.

**Supplementary Fig 7.**
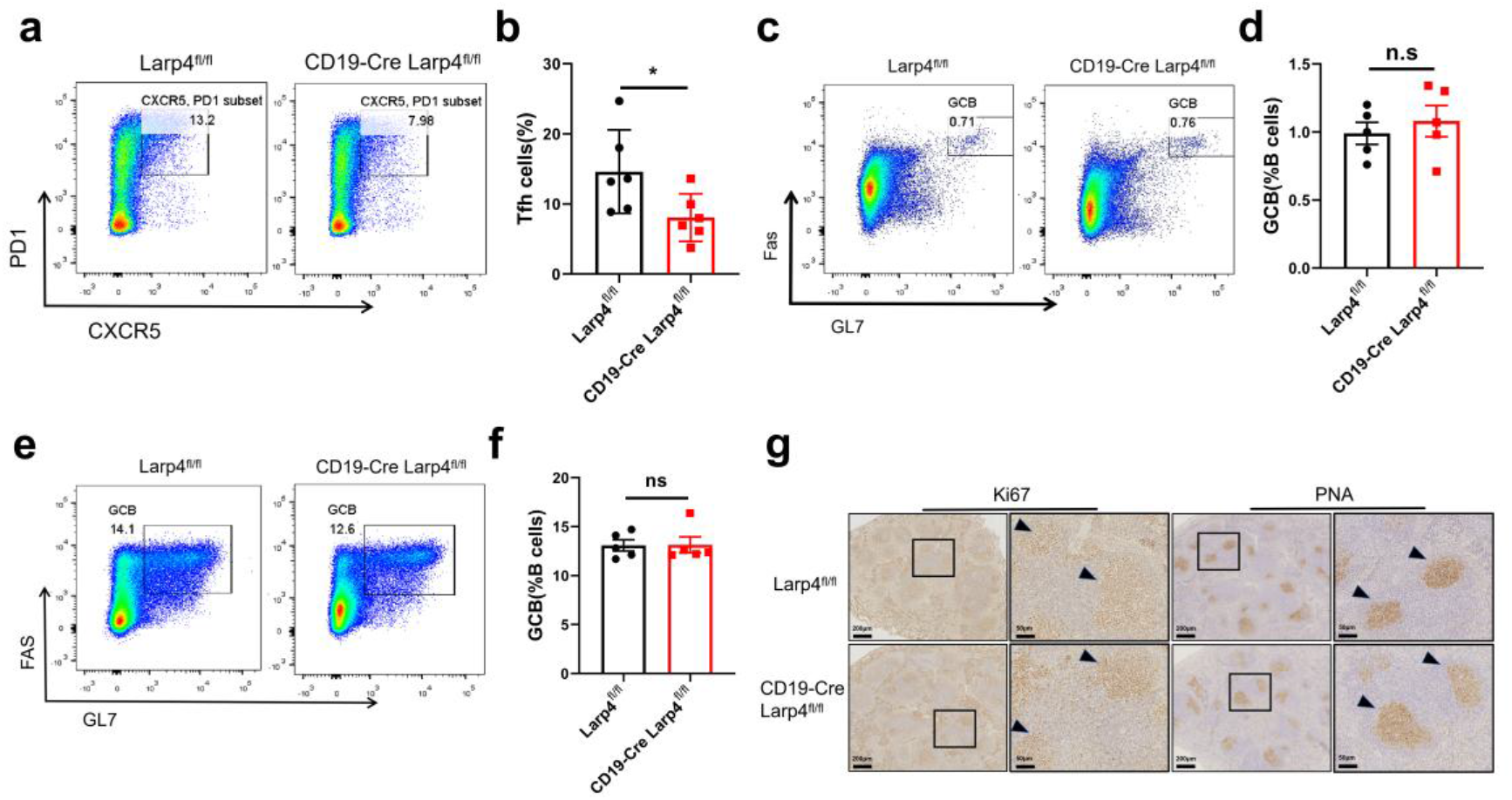
LARP4 deficiency preserves GCB cell formation in multiple immunization models. **a,b.** Flow cytometry analysis of splenic Tfh cells from Larp4^fl/fl^ and B-Larp4 CKO mice in the bm12-induced SLE model**(a)** and the percentage of Tfh cells is presented **(b)**. **c,d.** Flow cytometry analysis of GCBC in the spleens of Larp4^fl/fl^ and B-Larp4 CKO mice at day 28 with NP_33_-KLH immunization **(c)** and the percentage of plasma cells is presented **(d)**. **e,f.** Flow cytometry analysis of GCBC in the spleens of Larp4^fl/fl^ and B-Larp4 CKO mice at day 14 with SRBC immunization **(e)** and the percentage of plasma cells is presented **(f)**. **g**. Representative immunohistochemistry images of Ki67 (left) and PNA (right) staining of splenic germinal centers from Larp4^fl/fl^ and B-Larp4 CKO mice after SRBC immunization. Scale bar, 250 μm and 50 μm.

**Supplementary Fig 8.**
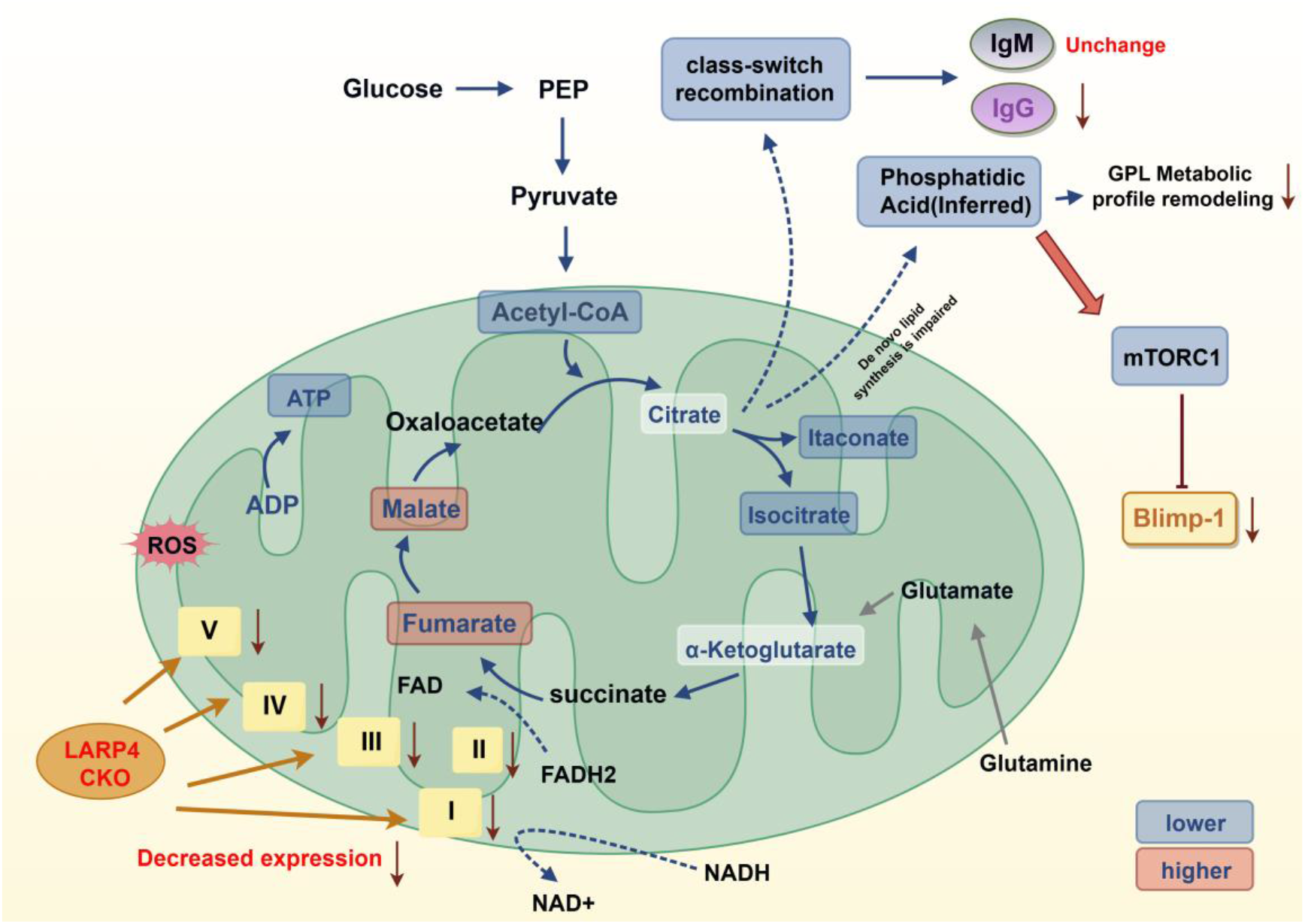
Schematic model illustrating the mechanism by which LARP4 deficiency blocks plasma cell differentiation. Schematic summary of the proposed mechanism. LARP4 deficiency impairs OXPHOS function and TCA cycle flux in activated B cells, leading to reduce de novo synthesis of phosphatidic acid (PA). Decreased PA levels attenuate mTORC1 activity, which in turn suppresses Blimp-1 upregulation, thereby selectively blocking plasma cell terminal differentiation. Related to Fig. 4-6.

**Supplementary Fig 9.**
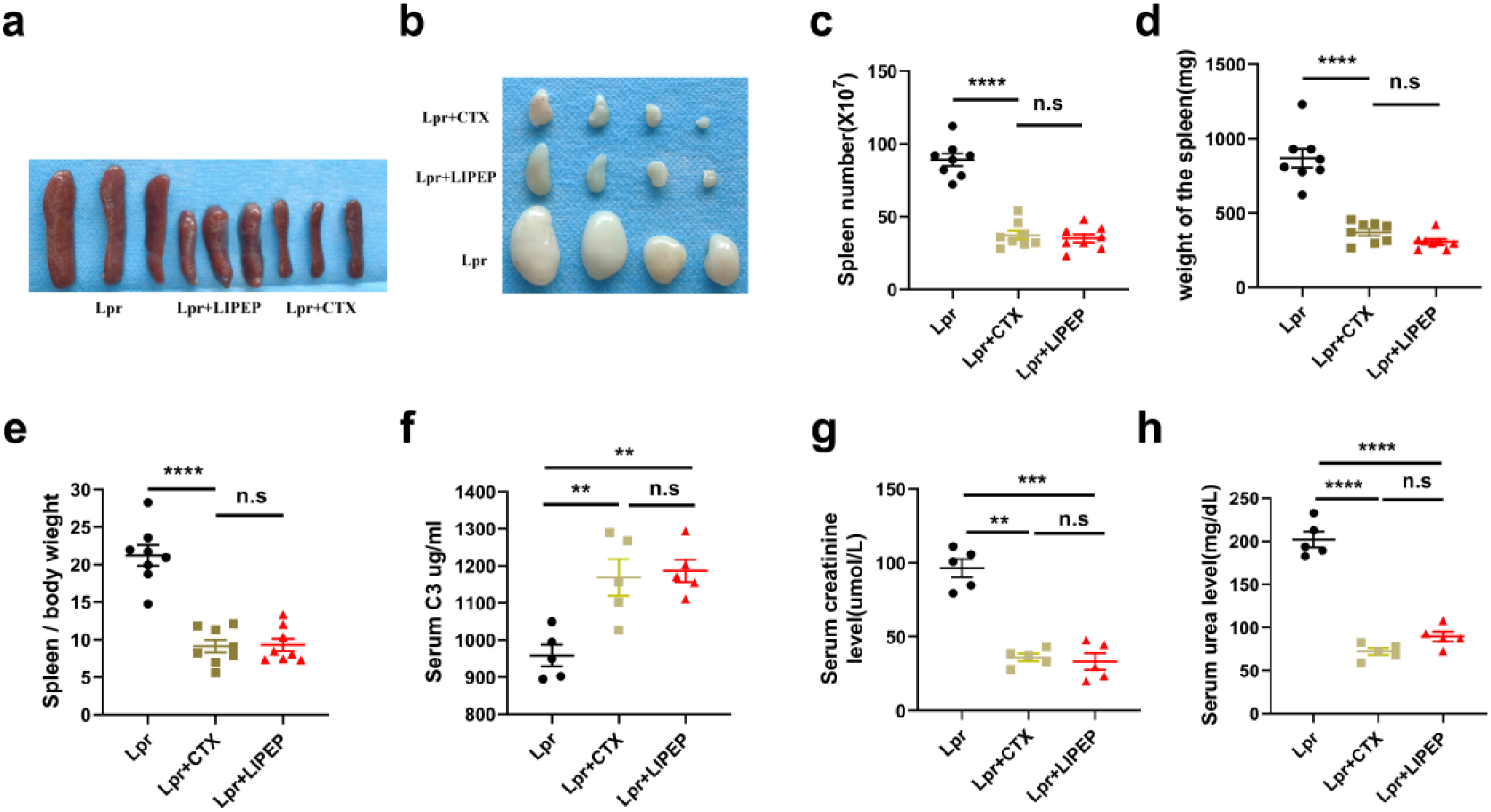
LIPEP treatment improves systemic and renal function parameters in MRL/lpr mice. **a-e.** Spleen size**(a)**, lymph node size**(b)**, spleen weight **(c)**, plenocyte counts **(d)** and spleen index (spleen weight/body weight, **e**) in DMSO, CTX, and LIPEP-treated MRL/lpr mice at 18 weeks of age. **f.** Serum complement C3 levels measured by ELISA. **g, h.** Serum creatinine (**g**) and blood urea nitrogen (BUN, **h**) levels in each group. n = 6-8 per group, *p < 0.05, **p < 0.01, ***p < 0.001,****p < 0.0001 versus Lpr. Data are presented as mean±SEM by one-way ANOVA with Tukey’s post-hoc test.

## Notes

### Competing Interest Statement

The authors have declared no competing interest.

